# Experimental Investigations of Impact Strength of Structural Wood Available in Northern Himalayas

**DOI:** 10.1101/2024.08.29.610261

**Authors:** Ummer Amin Sheikh, Azher Jameel, Majid Hameed Koul

**Affiliations:** Department of Mechanical Engineering, Islamic University of Science and Technology Awantipora, J&K, India; Department of Mechanical Engineering, National Institute of Technology Srinagar, J&K, India

**Keywords:** Structural wood, Impact strength, ASTM standards, Populus Nigra, Ulmus Wallichiana, Salix Alba Var, Juglus Riga, Cedrus Deodara, Genus Pinus

## Abstract

The current paper presents the detailed investigation of impact strength of different wood species available in the northern Himalayan region of India. Wood has always provided a potential structural material for constructing different engineering structures in these hilly areas. Although some studies have been performed to determine the strength of structural wood, but individual mechanical characterization of different wood species available in these areas needs immediate attention to design safe and reliable structures. Seven different wood species that have been traditionally used for constructional purpose in northern Himalayan region of India, have been selected for investigation. Test samples of same species have been collected from different regions of this area for the purpose of experimentation. The test samples selected for experimentation were defect free and were developed in accordance with ASTM: D-143 standards. In the current work, wood samples have been seasoned naturally and it has been made sure that the moisture content of the test specimens remains below the fiber saturation point. Due to the anisotropic behavior of wood, the impact strength has been investigated along both principal loading direction i.e. parallel and perpendicular to grains.

## Introduction

Wood is an important and potential structural material that has been traditionally used for constructing different structures in the northern Himalayan region of India including the state of Jammu and Kashmir. This is particularly due to its easy availability and problems faced in transporting material and equipment to these hilly areas. Structural wood is a natural polymeric material derived from the stems and branches of trees. The safety and reliability of wooden engineering structures primarily depends on the design data available for the structural wood available in that particular region. Although mechanical characterization of wood has been carried out to determine various strength related properties, but individual studies have not been performed on the wood species available in the northern Himalayan region of India. Therefore, determination of various mechanical properties of structural wood available in this area deserves special focus to ensure the safety and reliability of engineering structures present in this area. Structural wood can be classified into soft wood and hardwood. Hardwood comes from broad-leaved trees, mostly deciduous which often shed their leaves at the end of their growing season. Soft wood comes from trees having needle like leaves and are evergreen trees. Softwoods are not generally durable unless protected by preservatives. Hard woods are slow growing which makes them more expensive and are of high density and strength than soft woods. There is less dependent on preservatives compared to softwood.

Wood is anisotropic in nature and has property variations along different directions. It is a naturally occurring polymeric material that is highly valued and has a variety of applications due to its texture, colour, density and strength [1]. Wood is the third largest building material used for construction. It has been used by humans for thousands of years for various purposes, including construction in hilly and mountainous areas, fuel, furniture making and paper production [2]. Wooden structures are highly durable when properly treated, detailed and built, which can be seen in many historic buildings all around the world. Structural wood can easily be shaped and connected using nails, screws, bolts and dowels or adhesively bonded together [3, 4]. The biggest advantage of the wood is that it is the natural resource, makes it easily available and economically feasible and provides better insulation from cold. Many ancient houses and bridges built with timber can be seen even today. It’s a naturally occurring, renewable plant-based cellular resource with exceptional strength-to-weight ratios and distinctive structural qualities that make it suitable for wide range of applications [5]. Throughout history, various strengthening procedures have been proposed to improve the mechanical qualities of raw wood and wood-based products [6]. The mechanical characteristics of wood are related to its ability to withstand loads and include resistance to external stresses such cleavage, bending, shear, tension and compression [7–9].

The internal structure of any wood species depends on various factors such as age, growth conditions and environmental factors [10]. Tree trunk consists bark, cambium layer, xylem, heartwood, sapwood and pith, which can be seen in Figure 1. Bark is the outer most part which serves as a protective barrier against physical damage, pests and environmental factors and it transports sugars produced by the photosynthesis from the leaves to the rest of the tree.

**Figure 1:**
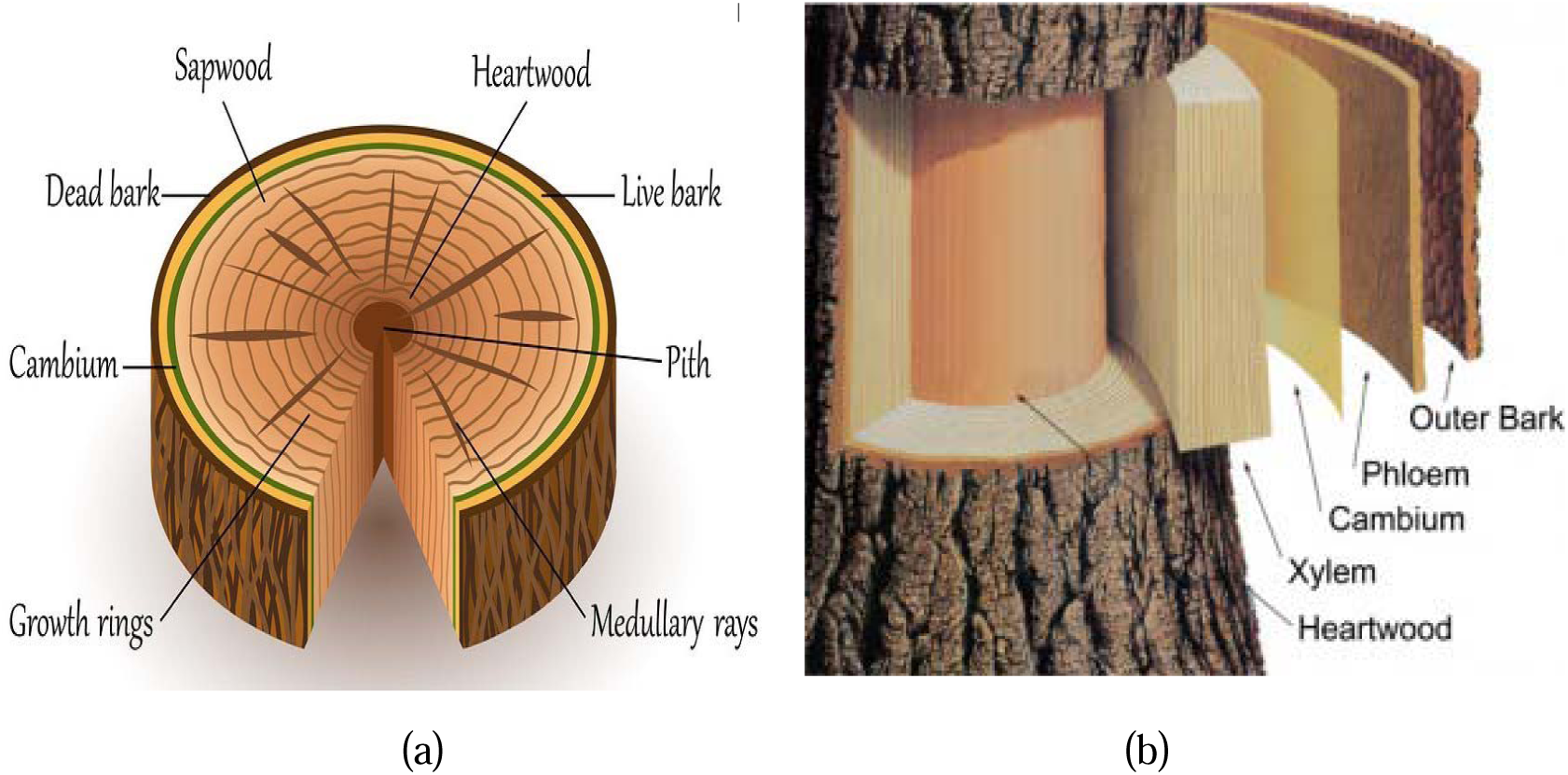
Internal structure of structural wood (a) Full cross-section (b) Different layers

Cambium layer is beneath the outer bark and is responsible for the trees growth in diameter. Xylem is inside the cambium layer and transports water and nutrients from the roots to the leaves and provides structural support to the tree. Each year the tree forms a new layer of xylem, resulting in the formation of annual rings that can be seen in cross-section of the trunk. Heartwood is at the centre of the trunk which consists of old wood and is the dead part of the tree trunk. Sap wood is surrounding to heartwood and is responsible for conducting water and nutrients. Sapwood decreases in width as the tree gets older and the area remains constant with age [11]. Heartwood is typically darker in colour and denser than sapwood and provides structural support to the tree. Sap wood is usually lighter in colour than heart wood and has less durability due to higher moisture content. Pith is the centre of the tree trunk and is formed during the early stage of the growth and serves as storage organ for reserves of water and nutrients. The internal structure of structural properties plays an important role in determining its mechanical properties [12–14]. Straight and parallel grained wood shows higher strength compared to irregular and sloping grains. This is because straight-grained wood distributes stresses more evenly along the length of the fibres, resulting in better structural integrity. Conversely, sloping or irregular grain may create weak points where the wood is more prone to splitting or breaking under stresses [15–17]. Straight-grained wood has a uniform, consistent appearance, making it desirable for applications where aesthetics are important, such as furniture making or woodworking crafts. It has been observed that the grain orientations have a significant impact on the mechanical properties of structural wood [18, 19]. In contrast, wood with irregular or sloping grain may exhibit unique patterns, which can be either desirable or undesirable depending on the specific application and personal preference. Wood with straight grain is generally easier to work with using hand or power tools whereas wood with irregular or sloping grains may be more challenging to work with, as the grain direction may change abruptly, leading to tear-out or splintering during cutting, planning, or sanding operations [20]. Wood with straight grain tends to be more stable and less prone to warping, twisting or cupping compared to wood with irregular or sloping grain. This is because straight-grained wood expands and contracts more evenly across the grains during changes in humidity which minimizes distortion. The mechanical properties of wood are mainly affected by its moisture content and grain direction, as it is an anisotropic bio-porous material [21–27]. The mechanical characteristics of wood define its suitability and capacity to withstand applied or external forces that could cause it to deform in any manner.

One of the important parameters which affect the mechanical properties of structural wood is its moisture content. In general, higher moisture content presents lower strength and lower moisture content gives higher strength to structural wood [28–31]. The moisture content present in structural wood should be below the fibre saturation point, for which seasoning of wood is required. The seasoning of timber refers to the process of drying wood to reduce its moisture content to a suitable level for its intended use. Properly seasoned wood is essential for ensuring stability, strength and durability in woodworking and construction projects. Seasoning can be either by natural or artificial seasoning. Immediately after harvesting, green wood typically contains a high moisture content in the range of 30% to over 200% of its dry weight, depending on the species and conditions [32]. Air drying is the most common method of seasoning timber. The rate of drying depends on factors such as ambient humidity, temperature, airflow and wood species [33–35]. Seasoning of wood must be carefully performed to minimize the occurrence of drying defects such as cups crooks, bows, top cracks and twists [36–38]. The ideal moisture content for seasoned timber varies depending on the intended use and environmental conditions and it typically ranges from 6% to 12%. Seasoning reduces the risk of shrinkage, warping, splitting, and decay that can occur when green or poorly dried wood is used. Well-seasoned timber is also easier to work with, as it is less prone to movement and distortion during machining, assembly, and finishing processes [39].

The current work reports the detailed investigation of impact strength of different wood species available in the northern Himalayan region of India including the state of Jammu and Kashmir. Seven different wood species that have been traditionally used for constructional purpose in northern Himalayan region of India, have been selected for investigation. These potential sources of structural wood include Populus Nigra, Ulmus Wallichiana, Salix Alba Var, Juglus Riga, Cedrus Deodara and Genus Pinus, as shown in Table 1. Due to the anisotropic behavior of wood, the impact strength has been investigated along both principal loading direction i.e. parallel and perpendicular to grains. The test samples selected for experimentation are defect free and have been developed in accordance with ASTM: D-143 standards. Natural air drying procedure has been used to reduce the moisture content of test samples below the fiber saturation point.

**Table 1:** Types of wood species found in northern Himalayan region (India)

| S.No | Botanical name | Local name | Family | Applications |
| --- | --- | --- | --- | --- |

**Table 1:** Types of wood species found in northern Himalayan region (India)
|  |  |  |  |  |
| --- | --- | --- | --- | --- |
| 1 | Populus<br>Ciliate | Poplar | Willow<br>Family<br>(Salicaceae) | Shutters, Pencils, trusses, Boxes<br>and crates, Framing, plywood and<br>furniture |
| 2 | Ulmus<br>Wallichiana | Elm | Ulmaceae<br>Family | Building trusses, roofs , interior<br>paneling, apple boxes, shutters,<br>furniture and flooring |
| 3 | Salix Alba Var | Willow | Willow<br>Family | Cricket Bats, Construction<br>Frames, packing frames, Kangris,<br>Baskets, paper industry, Chairs,<br>plywood and furniture |
| 4 | Juglus Riga | Walnut | Walnut<br>Family<br>(Juglandaceae) | Musical instruments, high quality<br>furniture, flooring , Baby toys,<br>interior paneling, carvings |
| 5 | Cedrus<br>Deodara | Deodar | Pine Family<br>(Pinaceae) | Beams, Floor-boards, window<br>frames Railway sleepers, posts<br>and door frames, furniture,<br>carvings, Interior paneling,<br>window frames, House Boats and<br>Bridges |
| 6 | Genus Pinus | Pine | Pine Family<br>(Pinaceae) | Beams , ply boards , ceilings,<br>doors, windows and platforms and<br>wall cladding, furniture, paneling,<br>window frames, roofing, window<br>frames, |
| 7 | Albies<br>Spectabilis | Silver Fir | Pine Family<br>(Pinaceae) | Beams, packing cases, doors,<br>windows drawers, and furniture |

## 2. Types of wood species and nomenclature

Different types of structural wood can be found in the northern Himalayan region of India. People living at higher attitudes mainly depend on wood for constructing different structures such as houses, bridges etc. Therefore, it is necessary to understand the type of wood available for construction in this region. Most commonly used structural wood comes from Poplar, Elm, Willow, Walnut, Deodar, Pine and Silver Fir. The details of different types of structural wood is given in this section.

### 2.1 Poplar (Populus Ciliate)

Poplar is locally known as ‘Fress’ for the people of Jammu and Kashmir. Its botanical name is Populus Ciliate. Popular wood is a hard wood and is whitish in colour as shown in figure 2. It is light in weight and can be easily worked. Poplar tree is straight grained fast growing long and straight round tree and is present in the bulk in the Himalayan region of India. It usually grows in low temperature areas where sandy clay is present. It grows along ponds, cannels or wet lands and needs less water content for its growth. The density of poplar wood is low and it dries very quickly. Structural wood obtained from poplar has a very low tendency to split when nailed and hence it has been a prime choice for structural constructions compared to willow, it is light in weight and has low shock resistance. Poplar wood has been traditionally used for making roofs of buildings, apple boxes and bridges.

**Figure 2:**
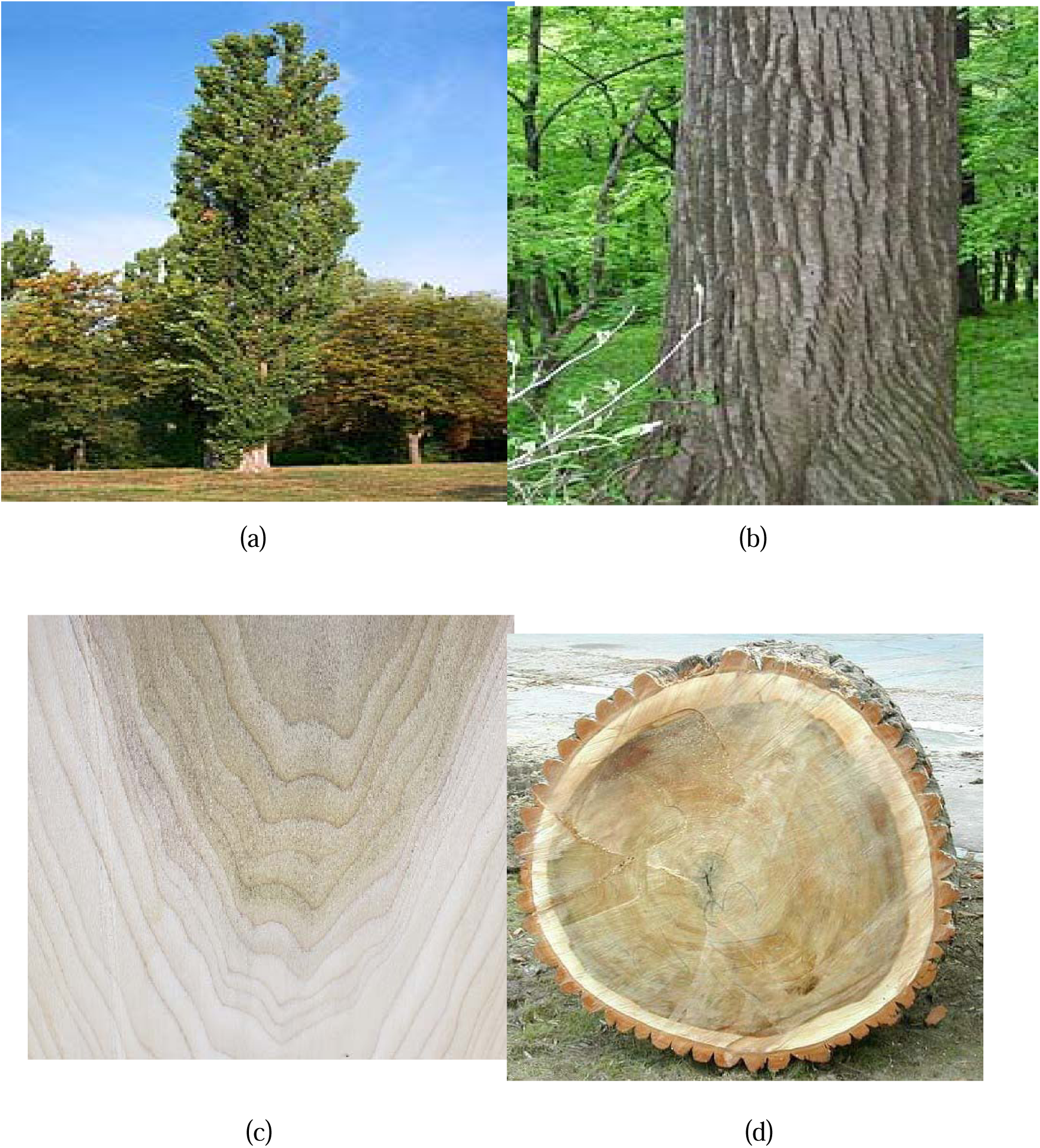
Details of poplar wood (a) poplar tree (b) poplar tree trunk (c) poplar wood with grains (d) cross-section of tree trunk

### 2.2 Elm (Ulmus Wallichiana)

Elm is locally known by the name ‘Brenn’ in Jammu and Kashmir. The botanical name of Elm is Ulmus Wallichiana. Elm wood falls under hardwood category and is softer than other counterparts. It is very heavy and dense and is chocolate red-brown in color, as shown in figure 3. It is considered as soft hard wood because of its strength, durability to hard wood but still softer than majority of hard woods. The grain pattern of Elm wood has pretty pattern. Elm wood is used for flooring, furniture, ceiling covers because of having high densities it is also used for panelling and for construction of musical instruments

**Figure 3:**
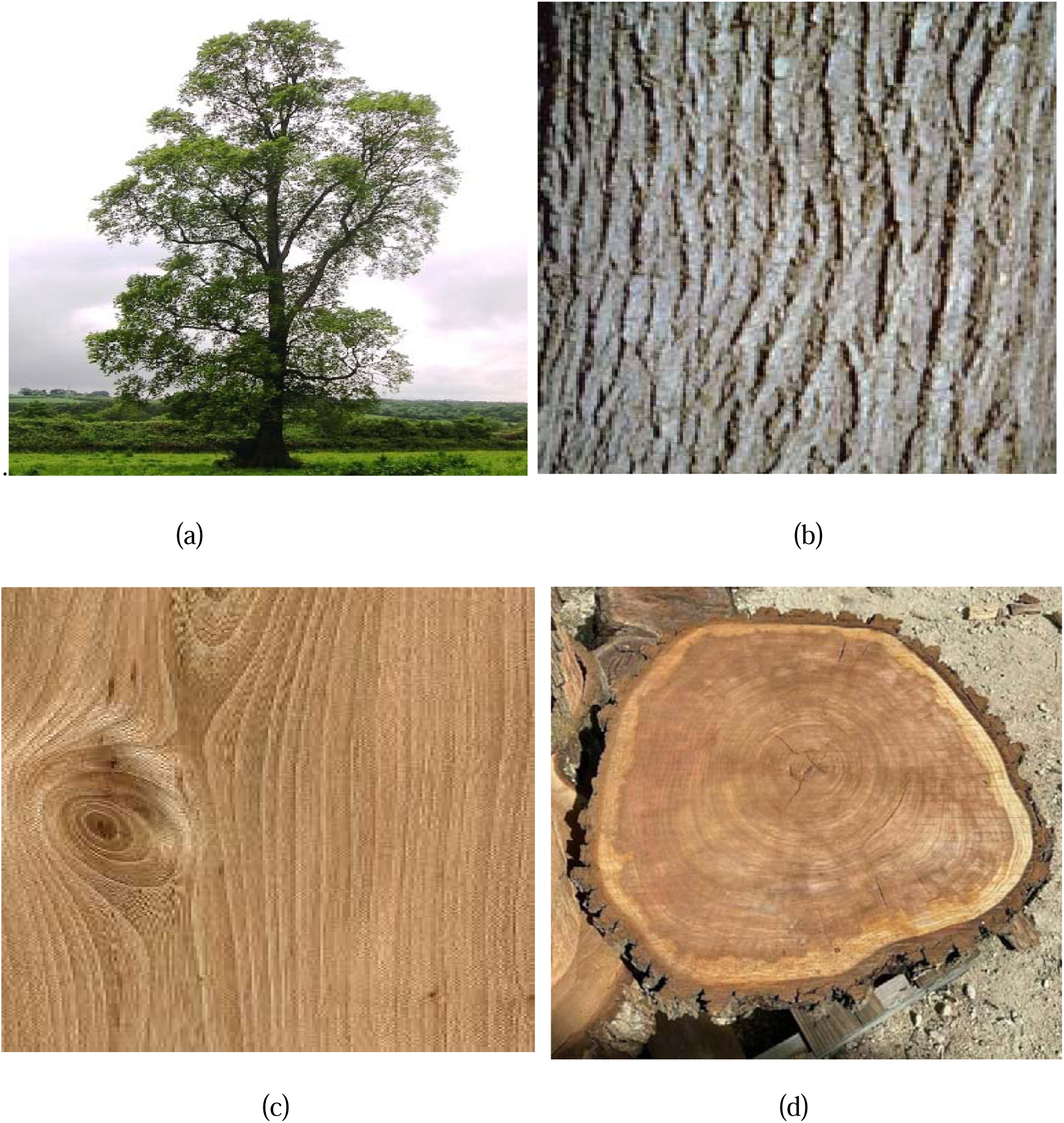
Details of Elm wood (a) Elm tree (b) Elm tree trunk (c) Elm wood with grains (d) Cross-section of tree trunk

### 2.3 Willow (Salix Alba Var)

The common name of willow in local language of Jammu and Kashmir is ‘veer’ and it is mostly found near the streams, rivers and brooks. The botanical name of willow is Salix Alba Var. Willow wood is straight grained hard wood but light in weight and is soft with good shock resistant can be easily worked. The willow tree, its bark and grain structure are shown in figure 4.Generally willow wood can be calorized into two type’s namely English willow with white leaves and Kashmiri willow with green leaves. Traditionally willow has been used for manufacturing bats and is a great choice to make cricket bats because of is high impact strength, light weight and good shock resistance. Small branches of this wood are used for making baskets, furniture, carvings and crates.

**Figure 4:**
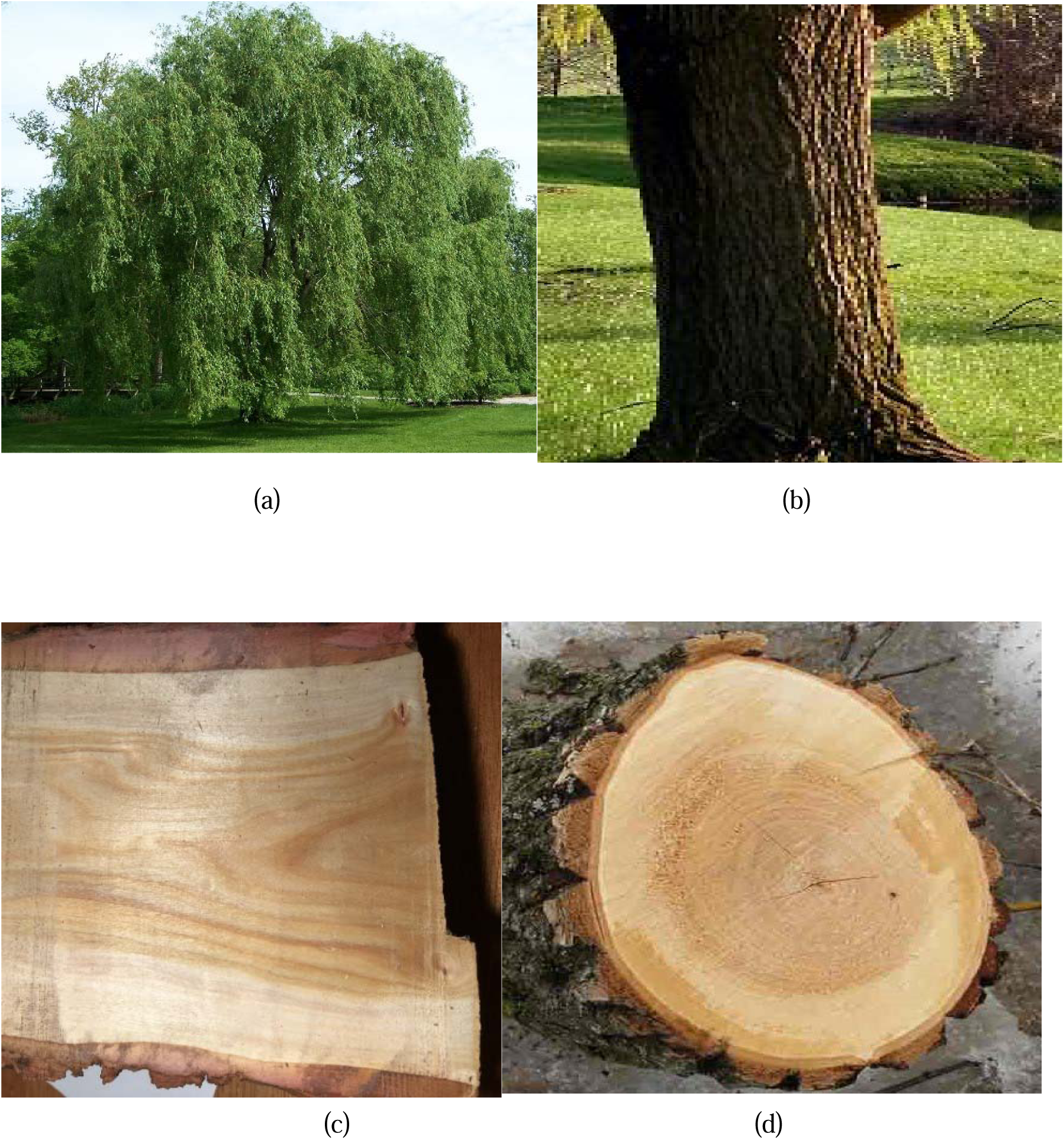
Details of Willow wood (a) willow tree (b) willow tree trunk (c) willow wood with grains (d) cross-section of tree trunk

### 2.4 Walnut (Juglus Riga)

Walnut tree is known by the local name ‘Doon Kull’ in Jammu and Kashmir and its botanical name is Juglus Riga. Walnut is tough hard wood and having low stiffness and moderate crushing and bending strength. The walnut wood is dark in color and has very pretty grain patterns that makes it highly priced. Walnut wood has a straight grain that can sometimes be wavy or curly adding to its visual appeal, as shown in figure 5. Walnut wood exhibits good dimensional stability and it is less prone to warping, shrinking, or swelling due to changes in moisture content. It is very strong and dense wood and is used for making furniture and wall paneling, flooring and ceiling houses. Walnut wood gives an aesthetic look to the structures because of its color and grain pattern. Walnut wood is widely used in crafting high-quality furniture. It also polishes well and takes finishes, stains, and oils beautifully. It resists decay and can withstand wear, making it suitable for both functional and decorative purposes.

**Figure 5:**
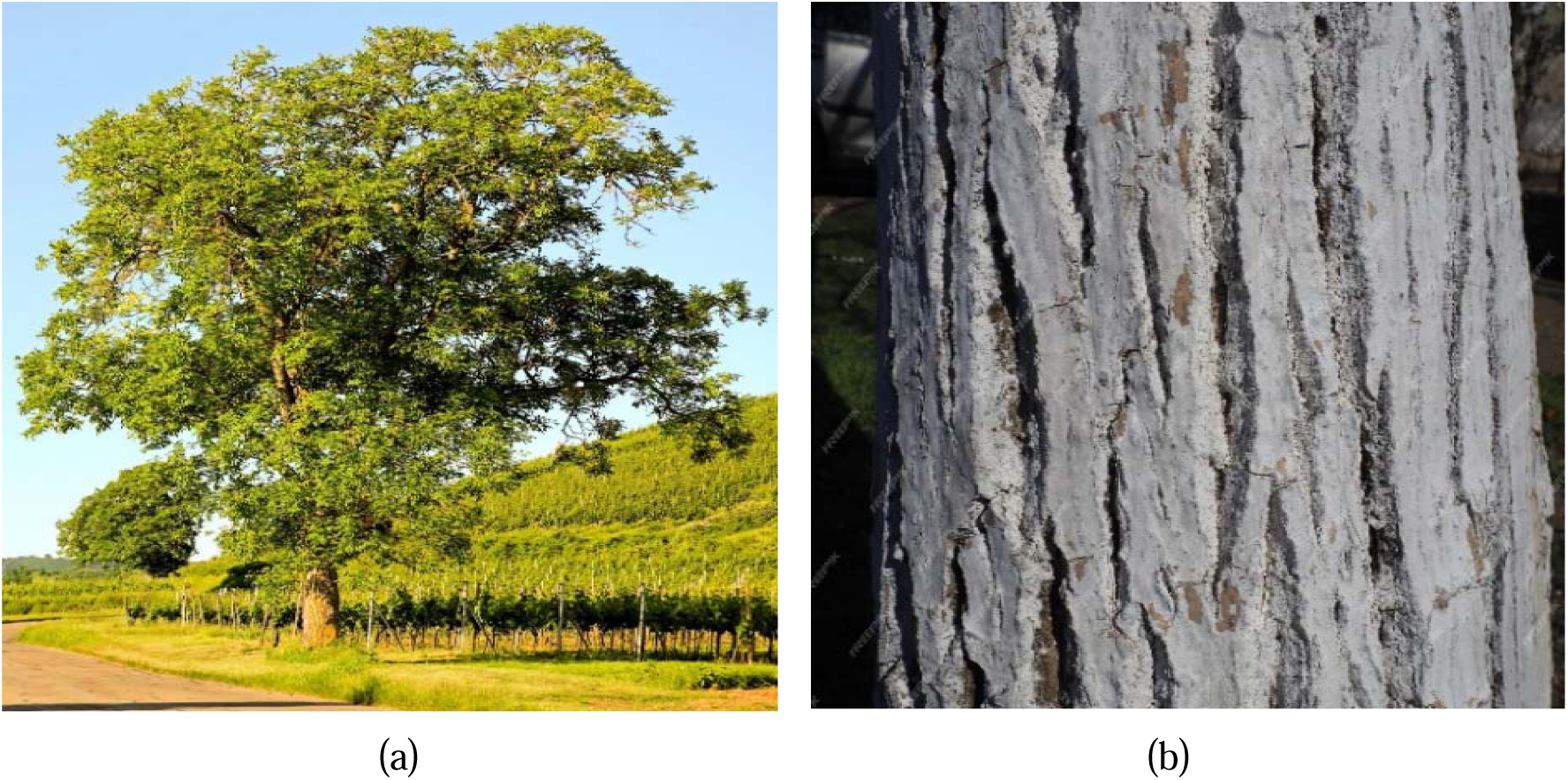
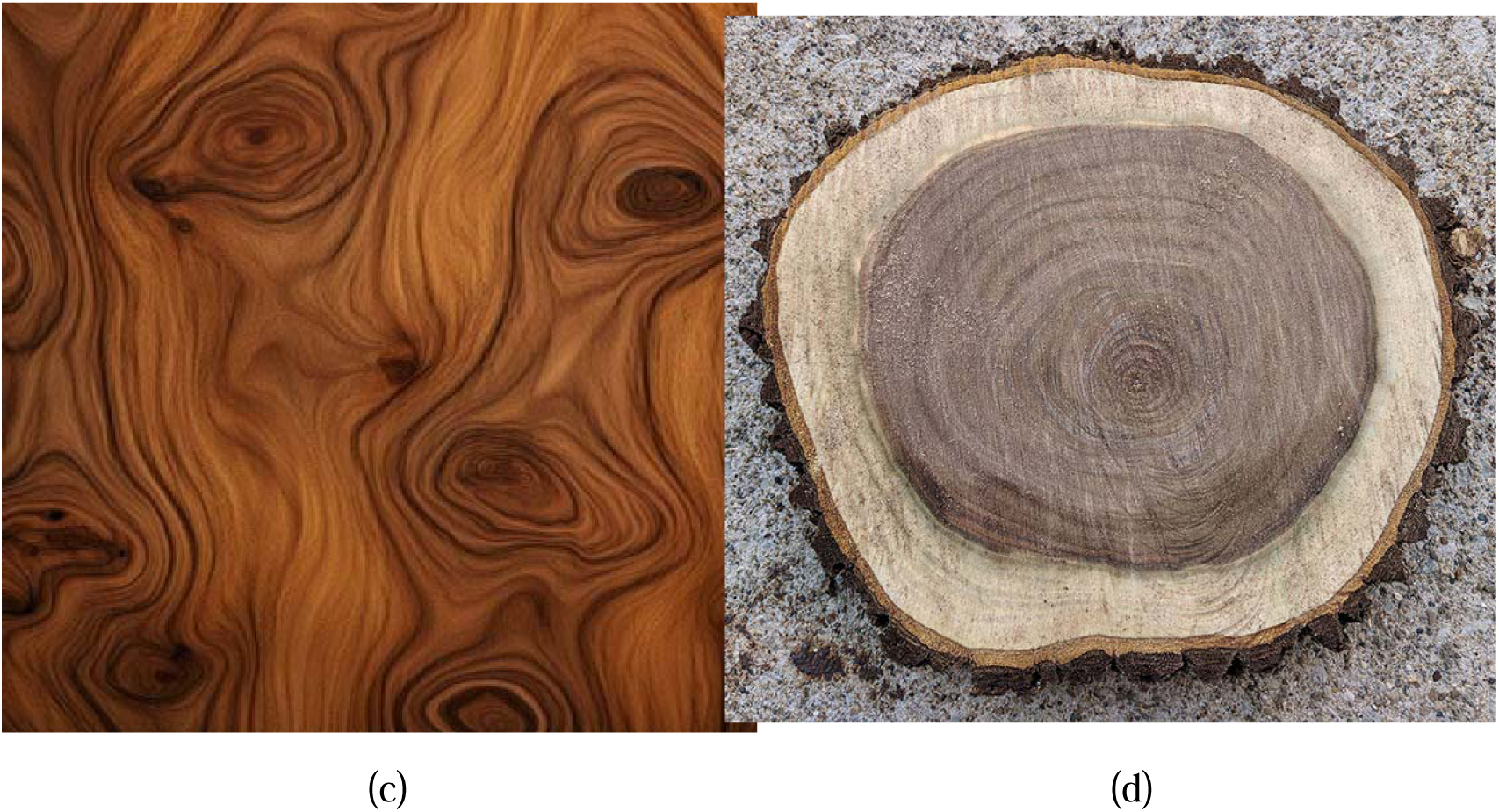
Details of Walnut wood (a) Walnut tree (b) Walnut tree trunk (c) Walnut wood with grains (d) cross-section of walnut tree trunk

### 2.5 Deodar (Cedrus Deodara)

Deodar is commonly known as Divdhor to the local population and its botanical name is Cedrus Deodara. Deodar wood is known for having a unique look and having straight long grains and its color ranges from light to reddish brown and the knots present in it enhance its aesthetic look. The constructional features of Deodar as shown ion figure 6. Deodar is soft wood and grows very tall and straight and is easily workable and machinable. Deodar has moderate strength and belongs to evergreen family having good smell. Deodar wood is known for its durability and resistance to decay. It possesses natural preservatives that make it resistant to insects and fungal attacks. Deodar has lower density as compared to other hardwoods but it provides the required strength and stability as required for constructional purposes. Deodar has always found to be a material of choice for machining doors, windows and as well as paneling. Because of its good resistance to moisture absorption, it has found extensive use in mainly house boats and bridges. It is suitable and safe for use in hot, moist and dry conditions. It is mostly used where the structure remains submerged in water for extended period of time and there is less effect on the life span of wood because of its water resistant nature.

**Figure 6:**
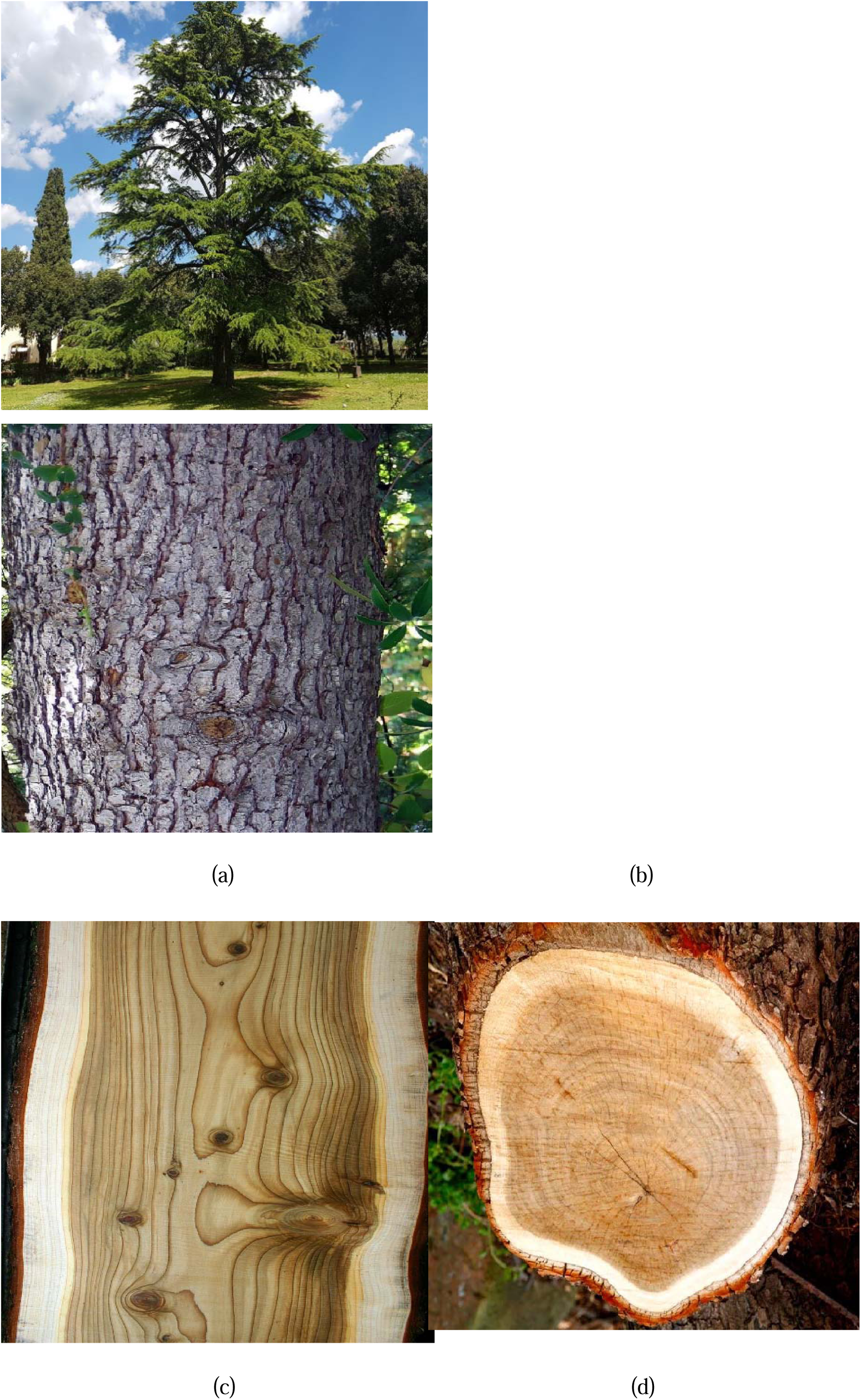
Details of Deodar wood (a) Deodar tree (b) Deodar tree trunk (c) Deodar wood with grains (d) cross-sectional of tree trunk

### 2.6 Blue Pine (Genus Pinus)

For local population pine is named as kaayur and its botanical name is Genus pine. It is found at higher attitudes and forest regions. Pine belongs to the category of soft wood and it comes from coniferous evergreen trees .it is very easy workable with using both hand and power tools. The problems of swelling or shrinking does not occur in pine wood. Pine typically has a straight grain pattern, although some species may have knots and irregulatries as shown in figure 7. It is lighter in weight and having excellent strength. Pine wood is used in construction of frames, sheathing, doors, windows and furniture. Pine wood is very much resistant to the damages caused by prolonged explosive to sunlight and hence it is used for constructing the structural parts that are exposed to sun light.

**Figure 7:**
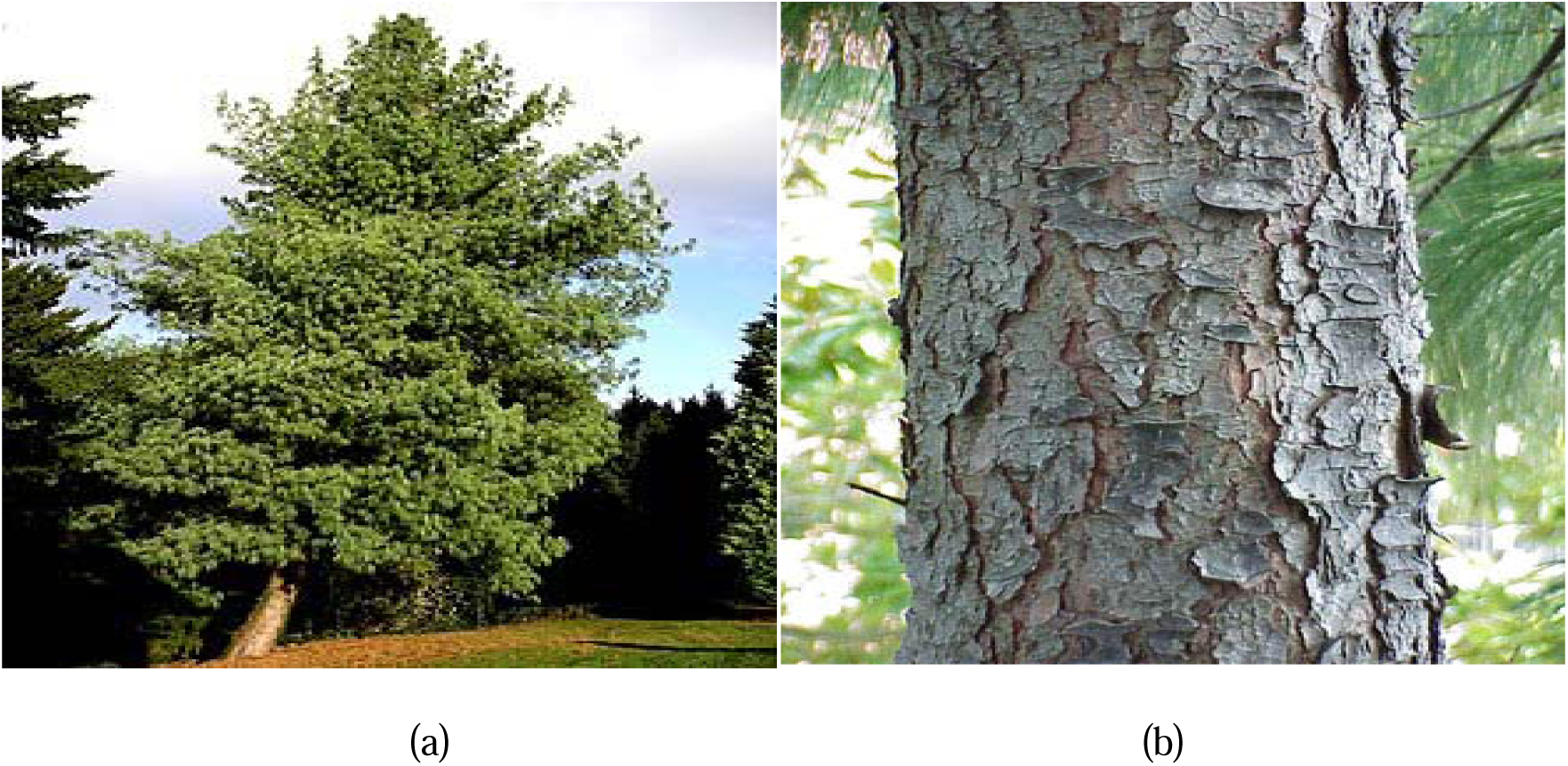
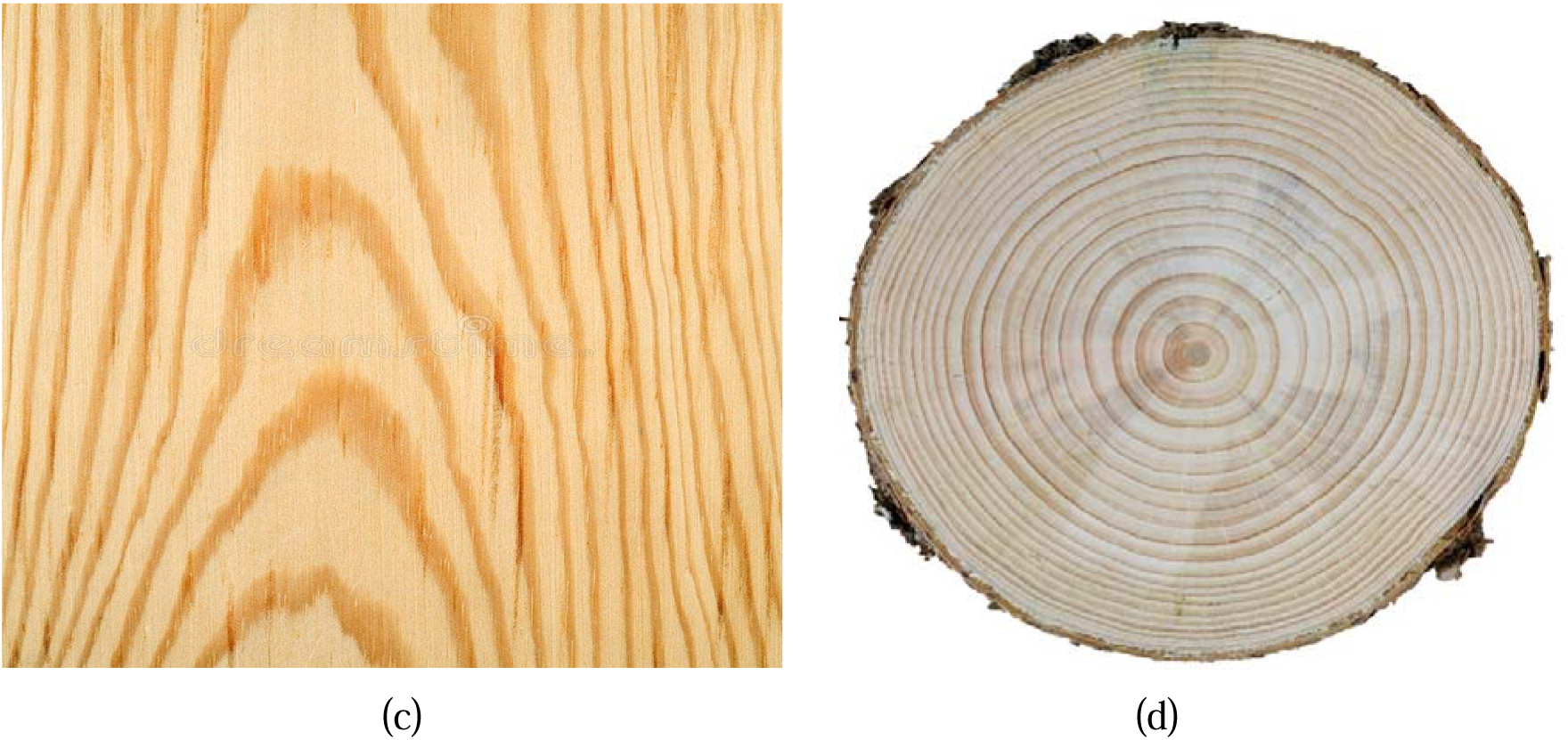
Details of Pine wood (a) Pine tree (b) Pine tree trunk (c) Pine wood with grains

### 2.7 Silver Fir (Albies Spectabilis)

The local name of silver fir in Kashmir is ‘Budul’ and its botanical name is Albies Spectabilis. It found at higher attitude locations. The silver fir is soft type of wood and clear straight grained and whitish in color as shown in figure 8. It is a preferred choice for local carpenters because it is easy to work, easily available and much cheaper as compared to pine and deodar. It is used to build doors and beams in wooden structures.

**Figure 8.**
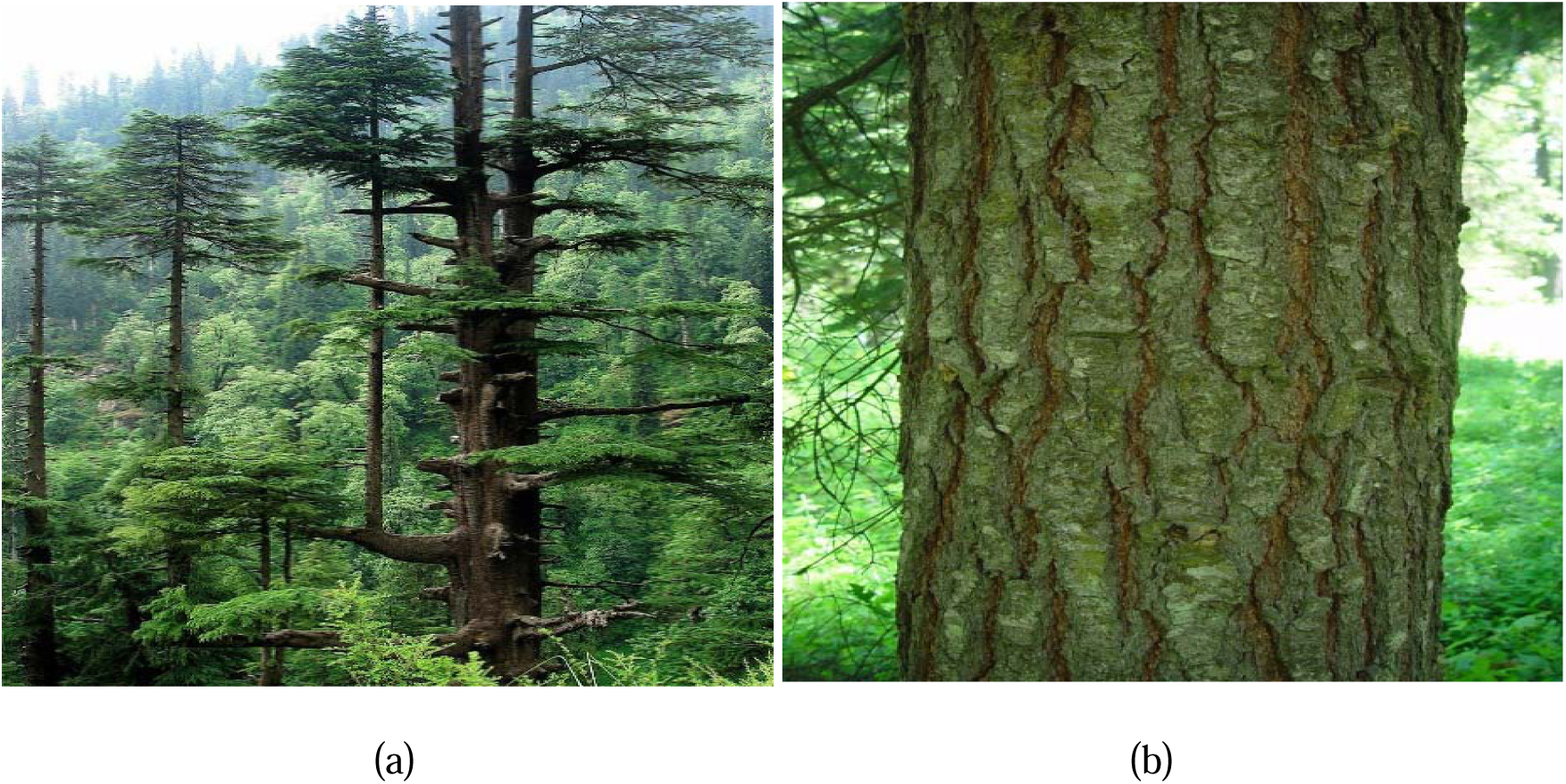
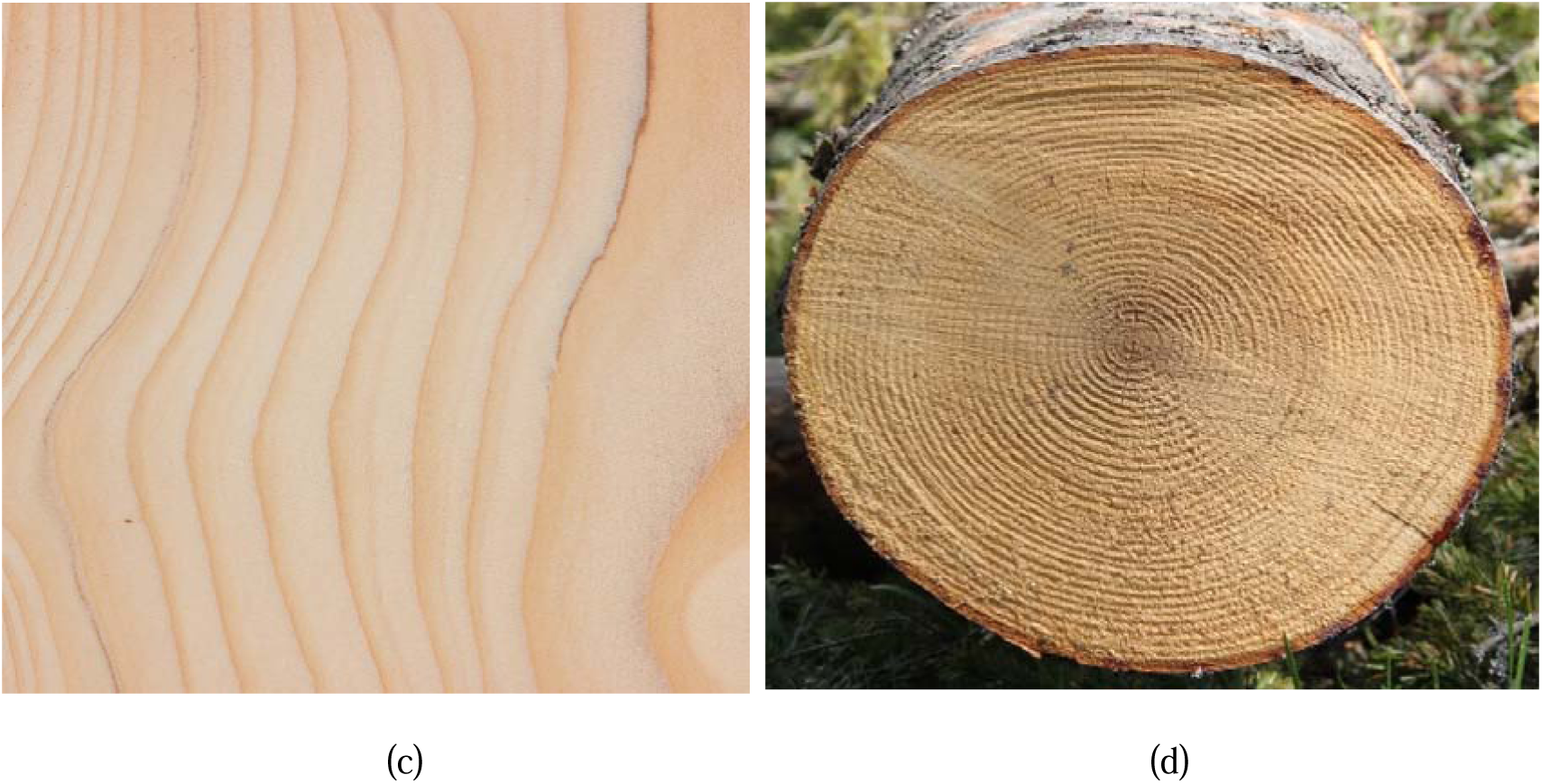
Details of Silver fir wood (a) Silver fir tree (b) Silver Fir tree trunk (c) Silver Fir tree wood with grains (d) cross-section of silver fir tree trunk

## 3. Materials and Methods

This section presents the details of the experimental methods adopted for estimating the impact strength of structural wood available in northern Himalayan region of India. Impact strength was calculated by performing the charpy and izod impact tests. Impact strength was calculated for structural wood in both the principle directions I.e. Parallel and perpendicular to grains. Seven different species of wood available in Jammu and Kashmir (India) were tested in both the directions parallel to grain and perpendicular to grain. From each type of wood ten specimens were selected for the purpose of experimentation. The test samples were collected from the different regions of Jammu and Kashmir (India). The specimens were obtained from different trees at the length of 1m to 3m from the bottom of the tree. The geometry of the specimens was selected according to applicable ASTM D codes which is ASTM D-143 for structural wood. The test specimens were seasoned by natural process without the use of artificial methods. It has been made sure that the moisture content of test samples remains below 12% at the time of experimentation. The moisture content of the wood specimens has kept below the Fiber Saturation Point. All the experiments performed on the state-of-act impact testing machine whose details are given in Figure 9 and Table 2.

**Figure 9:**
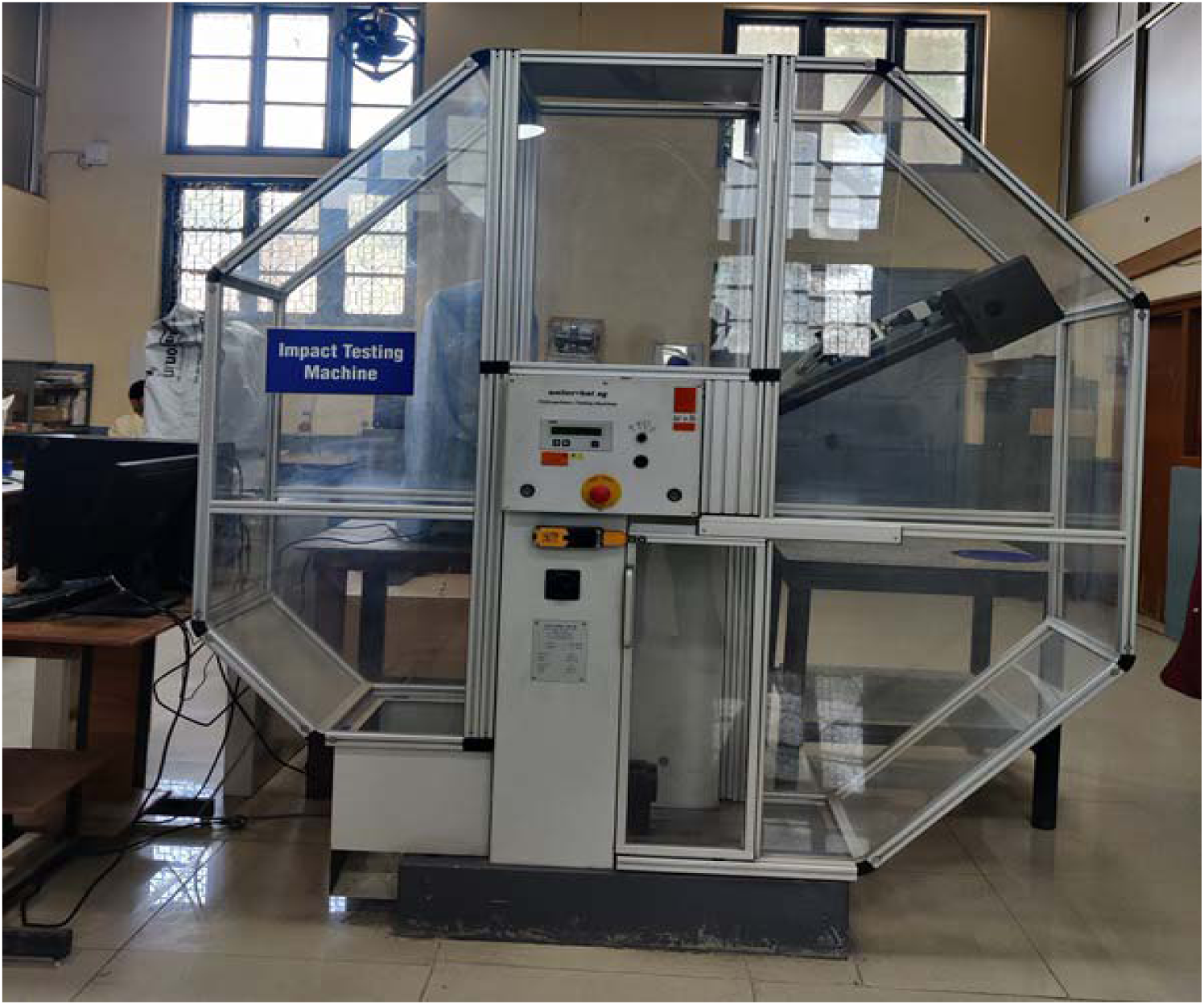
Impact testing machine used for experimentation

**Table 2:** Specifications of the Impact testing machine.

| Name of Machine | Make | Test Procedure | Pendulums |
| --- | --- | --- | --- |
| Impact Testing Machine | Walter + Bai | Charpy Test | Angle: 0° to 160°<br>Speed: 5.6 m/s<br>Length: 790 mm<br>Max. Energy: 450 J<br>Weight : 297 N |
|  |  | Izod Test | Angle: 0° to 160°<br>Speed: 5.6 m/s<br>Length: 790 mm<br>Max. Energy: 300 J<br>Weight: 195.7 N |

### 3.1 Standard procedure for Izod impact testing

The wooden specimens were prepared in the longitudinal direction for Izod impact testing parallel to the grain and the test specimens were prepared in the radial direction for parallel to grain. V-notch is used for conventional izod testing. The test specimen is typically a rectangular bar and the specimen dimensions are 75mm × 10mm × 10mm. The V-notch is given at the length of 28 mm with depth of 2 mm and subtended angle of 45 degree as shown in Figure 10. The dimensions, depth of notch, area of cross section and other geometrical features remains same for both parallel and perpendicular to grain tests. All test samples were naturally seasoned for three months so that the moisture content remains below the fiber saturation point. Natural seasoning occurred at room temperature where the relative humidity was equal to atmospheric humidity. Actual test specimens used in current work are shown in Figure 11.

**Figure 10.**
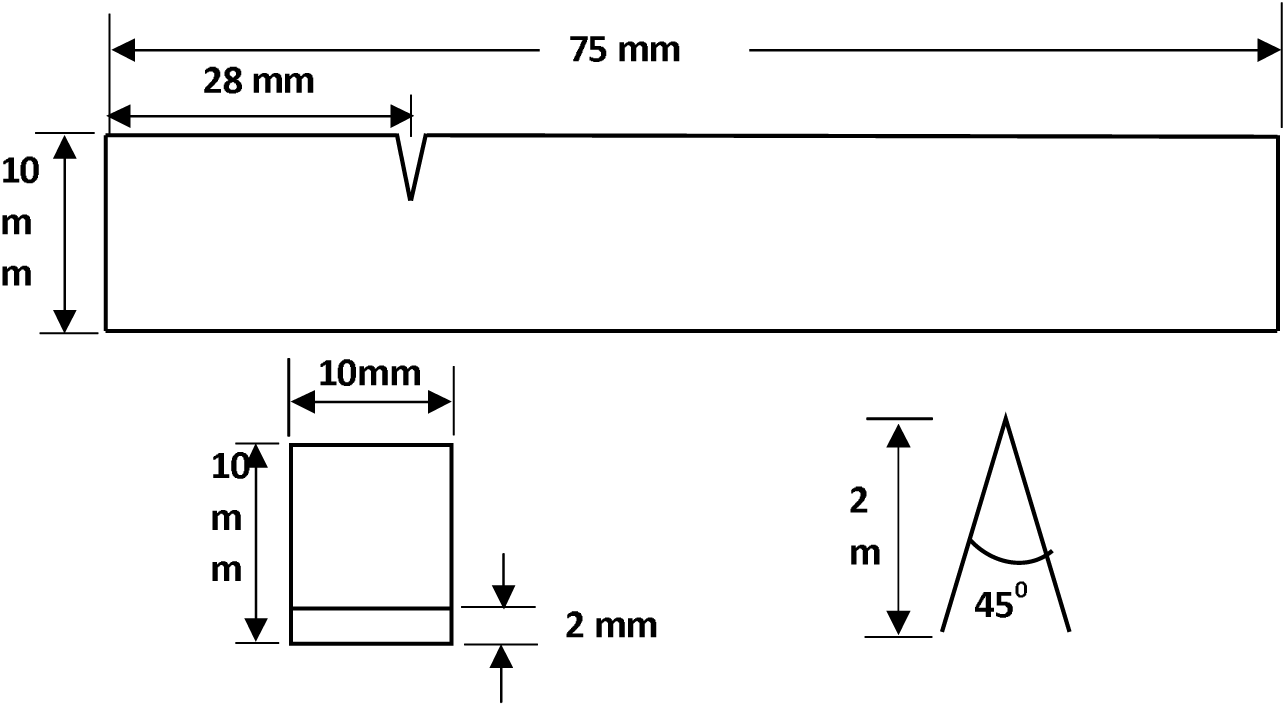
Standard test specimen for izod impact testing

**Figure 11.**
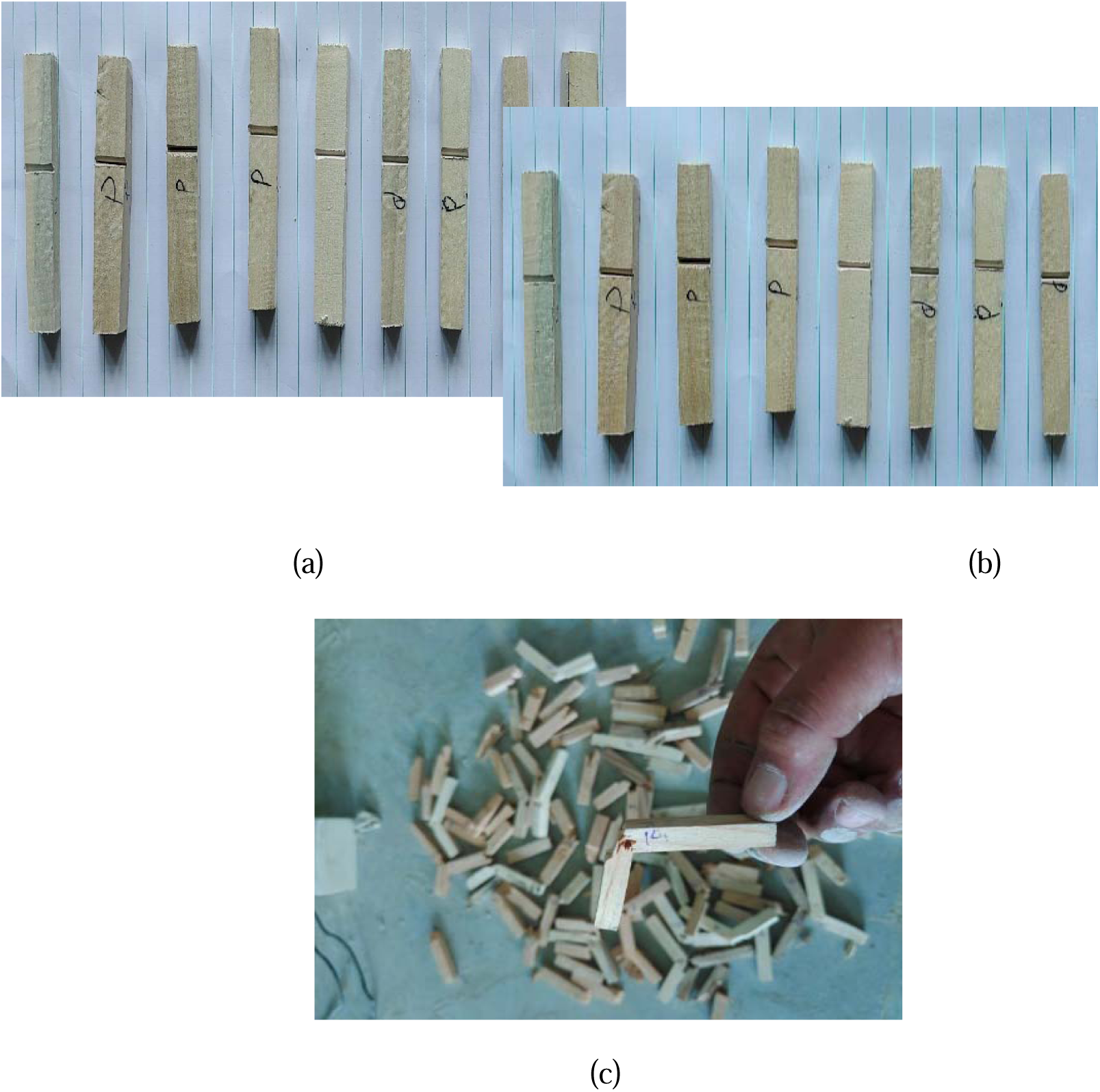
Test specimen for izod impact testing. (a) parallel to grain specimens (b) perpendicular to grain specimens (c) specimens after testing.

### 3.2 Standard procedure for Charpy impact testing

For Charpy test procedure, the test specimen is typically a rectangular bar and the specimen dimensions are 55mm × 10mm × 10mm. The V-notch is given at the center of the test specimen with depth of 2 mm and notch angle of 45 degree as shown in Figure 12. The dimensions, depth of notch, area of cross section and other geometrical features remains same for both parallel and perpendicular to grain tests. All test samples were naturally seasoned for three months so that the moisture content remains below the fiber saturation point. Natural seasoning occurred at room temperature where the relative humidity was equal to atmospheric humidity. Actual test specimens used in current work are shown in Figure 13.

**Figure 12.**
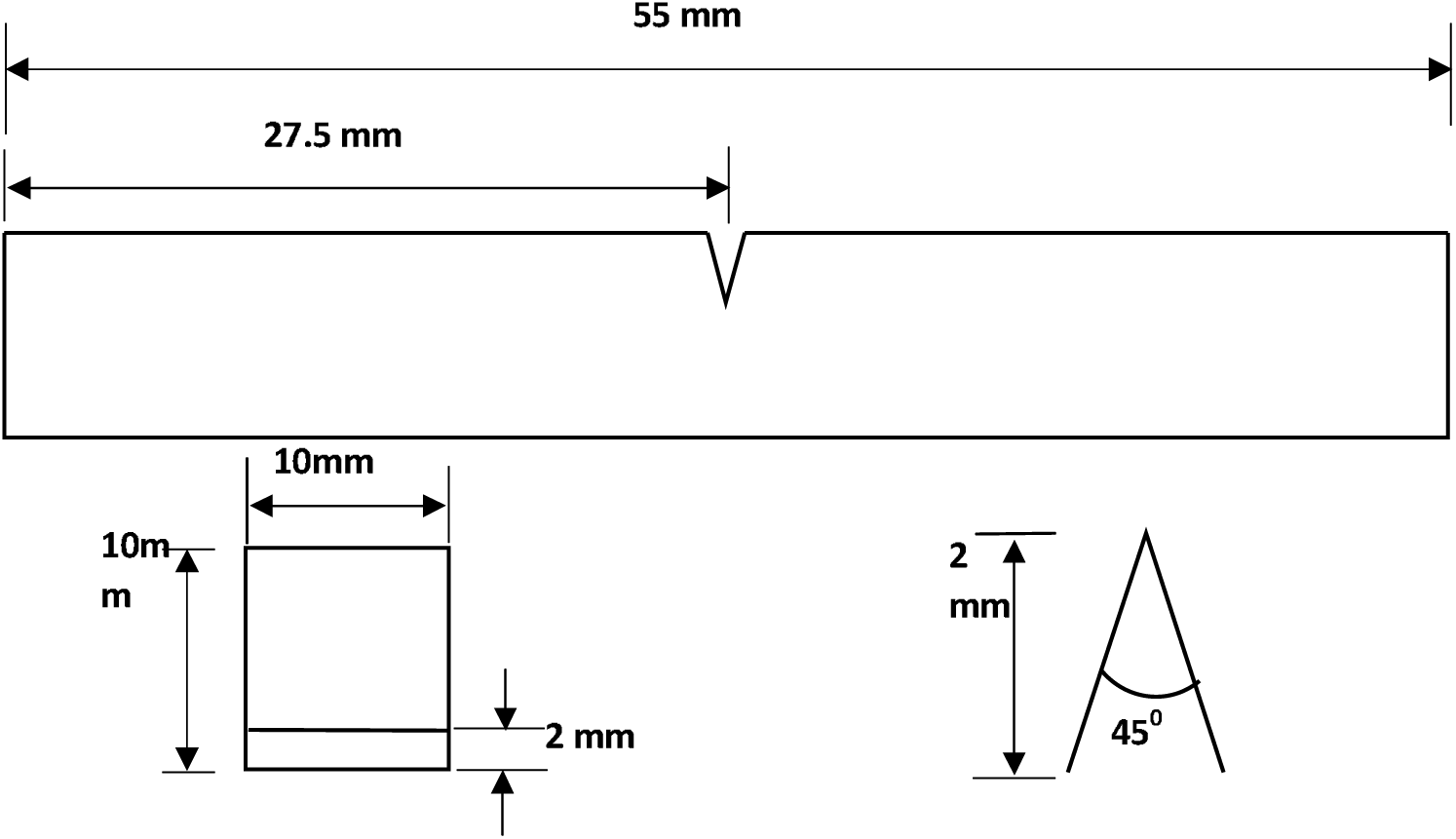
Standard Test Specimen for Charpy impact test

**Figure 13.**
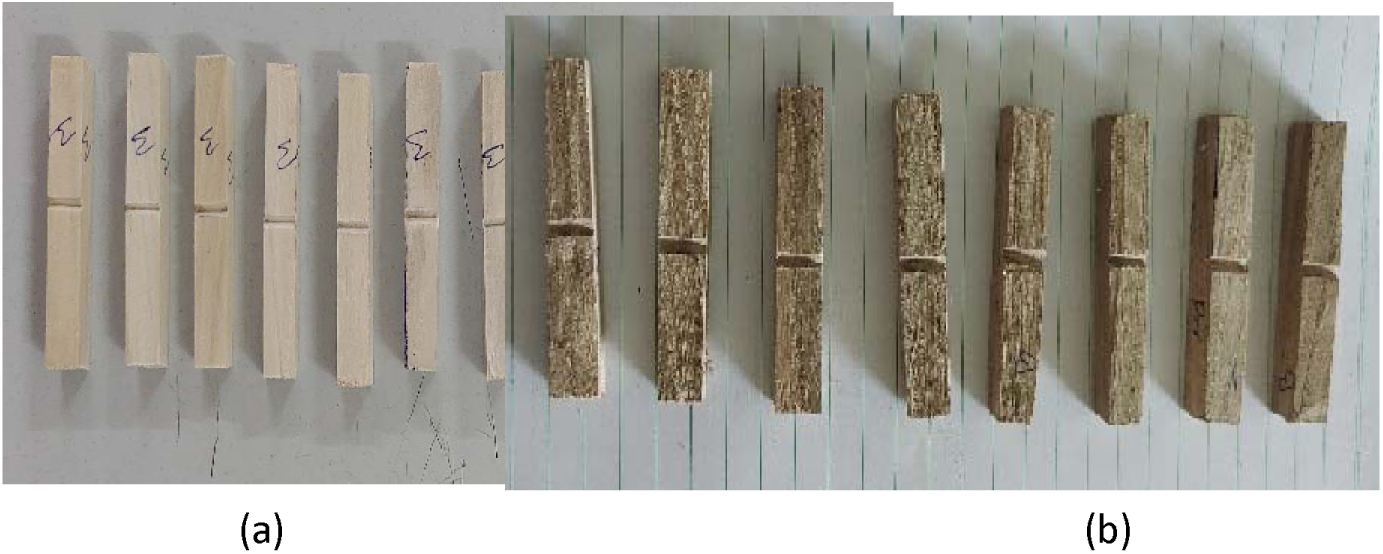
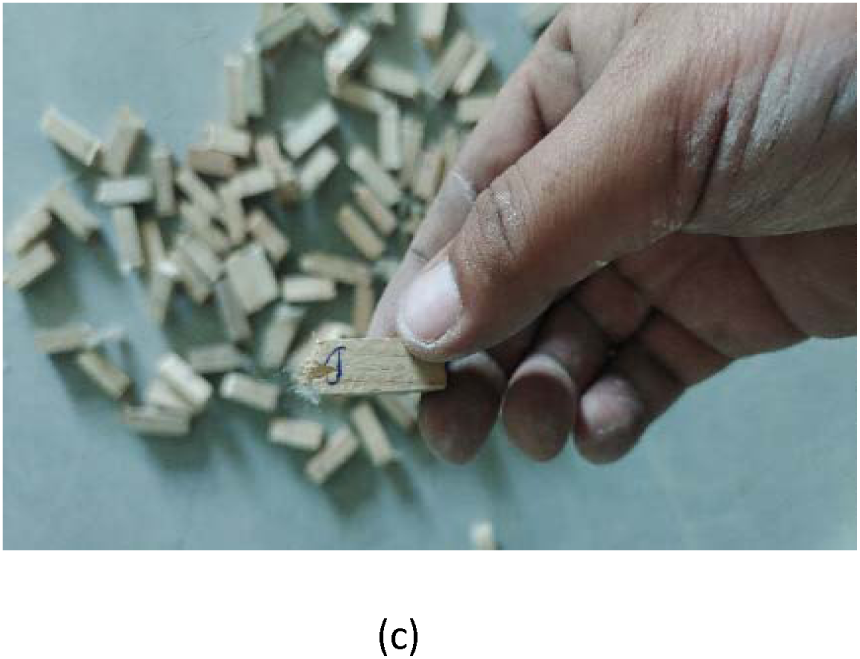
Test Specimen for Charpy Impact test (a) parallel to grain specimens (b) perpendicular to grain specimens (c) specimens after testing

## 4. Results and Discussions

This section presents the detailed investigations carried out on the impact strength of structural wood available in northern Himalayan regions of India. Test specimens have been obtained from seven different wood species that have been traditionally used for construction purposes in these areas. Such investigations are extremely important to ensure the safety and reliability of engineering structures constructed in these areas. The direction of loading and grain orientations have a significant influence on the impact strength of structural wood. In view of this, impact strength has been investigated in both principal directions i.e. parallel and perpendicular to grains. Both Izod and Charpy impact tests were used to obtain the impact strength of different wood species.

### 4.1 Impact Strength of Populus Nigra

Populus Nigra is locally known as “Poplar” to the people of Jammu and Kashmir and it has found extensive application in construction of different engineering structures. One of the potential applications of this wood species is the construction of frames for roof tops of buildings and bridges. In the current work, ten test samples were collected from Poplar trees available in different areas of northern Himalayan region. The test samples selected for experimentation were defect free and were developed in accordance with the ASTM: D-143 standards. The results obtained for the test samples, developed parallel to grains, using Izod impact testing are shown in Table 3. For the ten test samples, the minimum value of the impact strength was found to be 2.5 J/cm² whereas the maximum impact strength obtained was 3.5 J/cm². The average impact strength of Populus Nigra in the direction parallel to grains is found to be 2.79 J/cm^2^ by using Izod testing procedure. Impact strength has also been investigated in the direction perpendicular to grains. The impact strength of the test specimens, developed perpendicular to the grains, using Izod impact testing is shown in Table 4. As expected, the impact strength obtained for parallel to grain samples is much higher than those of perpendicular to grain test specimens. This is primarily due to the grain structures in two cases. For perpendicular to grain test specimens, the maximum impact strength was found to be 0.55 J/cm² whereas the minimum value was 0.23 J/cm². The average impact strength of Populus Nigra in the direction perpendicular to grains is estimated to be 0.37 J/cm^2^. From the experiments conducted in the current work, it was observed that Populus Nigra has lower impact strength as compared to other wood species available in this region, which will be discussed later.

**Table 3.** Impact Strength of Populus Nigra (Poplar) parallel to grains by Izod test procedure.

| Sample | Wood Species | Test Type | Impact strength |
| --- | --- | --- | --- |
| 1 | Populus Nigra<br>( Poplar) | IZOD | 2.50 J/cm<sup>2</sup> |
| 2 |  |  | 2.50 J/cm<sup>2</sup> |
| 3 |  |  | 3.50 J/cm<sup>2</sup> |
| 4 |  |  | 2.75 J/cm<sup>2</sup> |
| 5 |  |  | 2.50 J/cm<sup>2</sup> |
| 6 |  |  | 3.12 J/cm<sup>2</sup> |
| 7 |  |  | 2.75 J/cm<sup>2</sup> |
| 8 |  |  | 3.00 J/cm<sup>2</sup> |
| 9 |  |  | 2.75 J/cm<sup>2</sup> |
| 10 |  |  | 2.50 J/cm<sup>2</sup> |

**Table 4.** Impact Strength of Populus Nigra (Poplar) perpendicular to grains by Izod test procedure.

| Sample | Wood Species | Test Type | Impact strength |
| --- | --- | --- | --- |
| 1 |  |  | 0.30 J/cm<sup>2</sup> |
| 2 | Populus Nigra<br>(Poplar) | IZOD | 0.35 J/cm<sup>2</sup> |
| 3 |  |  | 0.45 J/cm<sup>2</sup> |
| 4 |  |  | 0.55 J/cm<sup>2</sup> |
| 5 |  |  | 0.25 J/cm<sup>2</sup> |
| 6 |  |  | 0.23 J/cm<sup>2</sup> |
| 7 |  |  | 0.34 J/cm<sup>2</sup> |
| 8 |  |  | 0.45 J/cm<sup>2</sup> |
| 9 |  |  | 0.40 J/cm<sup>2</sup> |
| 10 |  |  | 0.38 J/cm<sup>2</sup> |

Impact strength of Populus Nigra has also been evaluated by using the Charpy impact testing procedure. The impact strength of Populus Nigra, in the direction parallel to grains, by using the Charpy test procedure is given in Table 5. For the test samples obtained from different regions, the maximum impact strength parallel to grains was found to be 3.5 J/cm^2^ whereas the minimum impact strength along the same direction was observed as 2.5 J/cm^2^. The average impact strength of Populus Nigra, parallel to grains, was found out to be 2.84 J/cm^2^ by using the Charpy test procedure. Charpy impact tests were also performed on the samples obtained perpendicular to grains and the results obtained in the current work are shown in Table 6. For the test samples obtained perpendicular to the grains, the maximum impact strength was found to be 0.45 J/cm^2^ and the minimum value was 0.23 J/cm^2^. The average impact strength of Populus Nigra, perpendicular to grains, was found out to be 0.39 J/cm^2^.

**Table 5.** Impact Strength of Populus Nigra (Poplar) parallel to grains by Charpy test procedure.

| Sample | Wood Species | Test Type | Impact strength |
| --- | --- | --- | --- |
| 1 |  |  | 2.50 J/cm<sup>2</sup> |
| 2 |  |  | 3.50 J/cm<sup>2</sup> |
| 3 |  |  | 3.00 J/cm<sup>2</sup> |
| 4 |  |  | 2.75 J/cm<sup>2</sup> |
| 5 | Populus Nigra<br>(Poplar) | Charpy | 2.23 J/cm<sup>2</sup> |
| 6 |  |  | 3.13 J/cm<sup>2</sup> |
| 7 |  |  | 2.75 J/cm<sup>2</sup> |
| 8 |  |  | 3.25 J/cm<sup>2</sup> |
| 9 |  |  | 2.75 J/cm<sup>2</sup> |
| 10 |  |  | 2.50 J/cm<sup>2</sup> |

**Table 6.** Impact Strength of Populus Nigra (Poplar) perpendicular to grains by Charpy test procedure.

| Sample | Wood Species | Test Type | Impact strength |
| --- | --- | --- | --- |
| 1 | Populus Nigra<br>(poplar) | Charpy | 0.45 J/cm<sup>2</sup> |
| 2 |  |  | 0.40 J /cm<sup>2</sup> |
| 3 |  |  | 0.45 J/cm<sup>2</sup> |
| 4 |  |  | 0.36 J/cm<sup>2</sup> |
| 5 |  |  | 0.44 J/cm<sup>2</sup> |
| 6 |  |  | 0.23 J/cm<sup>2</sup> |
| 7 |  |  | 0.34 J/cm<sup>2</sup> |
| 8 |  |  | 0.45 J/cm<sup>2</sup> |
| 9 |  |  | 0.40 J/cm<sup>2</sup> |
| 10 |  |  | 0.38 J/cm<sup>2</sup> |

### 4.2 Impact Strength of Ulmus Wallichiana

Ulmus Wallichiana, locally known as Elm, is another important structural material in northern Himalayan region of India. Determination of impact strength of this wood species is very important to ensure the safety and reliability of the structures fabricated in these areas. For this purpose, ten defect free test samples were collected from different areas of region and these specimens were developed in accordance with the ASTM: D-143 standards. Impact strength of Ulmus Wallichiana parallel to grains using Izod impact testing procedure is shown in Table 7. The minimum value of the impact strength of this wood species was found to be 4.35 J/cm² whereas the maximum impact strength was 5.50 J/cm². The average impact strength of Ulmus Wallichiana in the direction parallel to grains is estimated to be 4.78 J/cm^2^ by using Izod testing procedure. Impact strength of this wood species was also evaluated in the direction perpendicular to grains. The impact strength perpendicular to the grains using Izod impact testing is shown in Table 8. It is observed that the impact strength of Elm parallel to grain is much higher than the strength obtained perpendicular to grains. The maximum impact strength perpendicular to grains was found to be 0.65 J/cm² whereas the minimum value was 0.45 J/cm². The average impact strength of Ulmus Wallichiana in the direction perpendicular to grains is estimated to be 0.53 J/cm^2^.

**Table 7.** Impact Strength of Ulmus Wallichiana (Elm) parallel to grains by Izod test procedure.

| Sample | Wood Species | Test Type | Impact strength |
| --- | --- | --- | --- |
| 1 | *Ulmus Wallichiana*<br>(Elm) | IZOD | 5.00 J/cm<sup>2</sup> |
| 2 |  |  | 4.50 J/cm<sup>2</sup> |
| 3 |  |  | 4.35 J/cm<sup>2</sup> |
| 4 |  |  | 4.50 J/cm<sup>2</sup> |
| 5 |  |  | 4.75 J/cm<sup>2</sup> |
| 6 |  |  | 4.50 J/cm<sup>2</sup> |
| 7 |  |  | 5.00 J/cm<sup>2</sup> |
| 8 |  |  | 5.18 J/cm<sup>2</sup> |
| 9 |  |  | 4.50 J/cm<sup>2</sup> |
| 10 |  |  | 5.50 J/cm<sup>2</sup> |

**Table 8.** Impact Strength of Ulmus Wallichiana (Elm) perpendicular to grains by Izod test procedure.

| Sample | Wood Species | Test Type | Impact strength |
| --- | --- | --- | --- |
| 1 |  |  | 0.45 J/cm<sup>2</sup> |
| 2 | Ulmus Wallichiana (Elm) | IZOD | 0.55 J/cm<sup>2</sup> |
| 3 |  |  | 0.65 J/cm<sup>2</sup> |
| 4 |  |  | 0.56 J/cm<sup>2</sup> |
| 5 |  |  | 0.45 J/cm<sup>2</sup> |
| 6 |  |  | 0.48 J/cm<sup>2</sup> |
| 7 |  |  | 0.61 J/cm<sup>2</sup> |
| 8 |  |  | 0.56 J/cm<sup>2</sup> |
| 9 |  |  | 0.48 J/cm<sup>2</sup> |
| 10 |  |  | 0.50 J/cm<sup>2</sup> |

In this study, impact strength of Ulmus Wallichiana has also been evaluated by using the Charpy impact testing procedure. The impact strength of Ulmus Wallichiana, in the direction parallel to grains, by using the Charpy test procedure is given in Table 9. The maximum impact strength parallel to grains was found to be 5.53 J/cm^2^ whereas the minimum impact strength along the same direction was observed as 4.3 J/cm^2^. The average impact strength of Ulmus Wallichiana parallel to grains has been evaluated to be 5.15 J/cm^2^ by using the Charpy test procedure. Charpy impact tests were also performed on the samples obtained perpendicular to grains and the results obtained in the current work are shown in Table 10. For the test samples obtained perpendicular to the grains, the maximum impact strength was found to be 0.56 J/cm^2^ and the minimum value was 0.43 J/cm^2^. The average impact strength of Ulmus Wallichiana, perpendicular to grains, was found out to be 0.49 J/cm^2^.

**Table 9.** Impact Strength of Ulmus Wallichiana (Elm) parallel to grains by Charpy test procedure.

| Sample | Wood Species | Test Type | Impact strength |
| --- | --- | --- | --- |
| 1 |  |  | 5.00 J/cm<sup>2</sup> |
| 2 |  |  | 5.53 J/cm<sup>2</sup> |
| 3 | Ulmus Wallichiana (Elm) | Charpy | 5.50 J/cm<sup>2</sup> |
| 4 |  |  | 5.00 J/cm<sup>2</sup> |
| 5 |  |  | 5.52 J/cm<sup>2</sup> |
| 6 |  |  | 4.30 J/cm<sup>2</sup> |
| 7 |  |  | 5.20 J/cm<sup>2</sup> |
| 8 |  |  | 5.50 J/cm<sup>2</sup> |
| 9 |  |  | 4.50 J/cm<sup>2</sup> |
| 10 |  |  | 5.50 J/cm<sup>2</sup> |

**Table 10.** Impact Strength of Ulmus Wallichiana (Elm) perpendicular to grains by Charpy test procedure.

| Sample | Wood Species | Test Type | Impact strength |
| --- | --- | --- | --- |
| 1 | Ulmus Wallichiana (Elm) | Charpy | 0.45 J/cm<sup>2</sup> |
| 2 |  |  | 0.44 J/cm<sup>2</sup> |
| 3 |  |  | 0.56 J/cm<sup>2</sup> |
| 4 |  |  | 0.56 J/cm<sup>2</sup> |
| 5 |  |  | 0.45 J/cm<sup>2</sup> |
| 6 |  |  | 0.43 J/cm<sup>2</sup> |
| 7 |  |  | 0.47 J/cm<sup>2</sup> |
| 8 |  |  | 0.56 J/cm<sup>2</sup> |
| 9 |  |  | 0.46 J/cm<sup>2</sup> |
| 10 |  |  | 0.50 J/cm<sup>2</sup> |

### 4.3 Impact Strength of Salix Alba Var

Salix Alba Var, locally known as Willow, is another important structural material that has been extensively used in the manufacture of cricket bats and other engineering structures. Same test procedures were adopted for determining the impact strength of willow as discussed in previous sections. Impact strength of Salix Alba Var parallel to grains using Izod impact testing procedure is shown in Table 11. The minimum value of the impact strength of this wood species was found to be 3.5 J/cm² whereas the maximum impact strength was 4.5 J/cm². The average impact strength of Salix Alba Var in the direction parallel to grains is estimated to be 4.00 J/cm^2^ by using Izod testing procedure. Impact strength has also been evaluated in the direction perpendicular to grains. The impact strength of this wood species perpendicular to the grains using Izod impact testing is shown in Table 12. The maximum impact strength perpendicular to grains was found to be 0.48 J/cm² whereas the minimum value was 0.32 J/cm². The average impact strength of Salix Alba Var in the direction perpendicular to grains is estimated to be 0.38 J/cm^2^.

**Table 11.** Impact Strength of Salix Alba Var parallel to grains by Izod test procedure.

| Sample | Wood Species | Test Type | Impact Strength |
| --- | --- | --- | --- |
| 1 | Salix Alba Var<br>(Willow) | IZOD | 4.00 J/cm<sup>2</sup> |
| 2 |  |  | 3.50 J/cm<sup>2</sup> |
| 3 |  |  | 4.50 J/cm<sup>2</sup> |
| 4 |  |  | 3.75 J/cm<sup>2</sup> |
| 5 |  |  | 4.00 J/cm<sup>2</sup> |
| 6 |  |  | 4.25 J/cm<sup>2</sup> |
| 7 |  |  | 3.50 J/cm<sup>2</sup> |
| 8 |  |  | 3.75 J/cm<sup>2</sup> |
| 9 |  |  | 4.50 J/cm<sup>2</sup> |
| 10 |  |  | 4.25 J/cm<sup>2</sup> |

**Table 12.** Impact Strength of Salix Alba Var perpendicular to grains by Izod test procedure.

| Sample | Wood Species | Test Type | Impact Strength |
| --- | --- | --- | --- |
| 1 | Salix Alba Var<br>(Willow) | IZOD | 0.34 J/cm<sup>2</sup> |
| 2 |  |  | 0.33 J/cm<sup>2</sup> |
| 3 |  |  | 0.32 J/cm<sup>2</sup> |
| 4 |  |  | 0.36 J/cm<sup>2</sup> |
| 5 |  |  | 0.36 J/cm<sup>2</sup> |
| 6 |  |  | 0.46 J/cm<sup>2</sup> |
| 7 |  |  | 0.40 J/cm<sup>2</sup> |
| 8 |  |  | 0.45 J/cm<sup>2</sup> |
| 9 |  |  | 0.48 J/cm<sup>2</sup> |
| 10 |  |  | 0.34 J/cm<sup>2</sup> |

In this study, impact strength of Salix Alba Var has also been evaluated by using the Charpy impact testing procedure. The impact strength of Salix Alba Var, in the direction parallel to grains, by using the Charpy test procedure is given in Table 13. The maximum impact strength parallel to grains was found to be 4.57 J/cm^2^ whereas the minimum impact strength along the same direction was observed as 3.35 J/cm^2^. The average impact strength of Salix Alba Var parallel to grains has been evaluated to be 3.91. J/cm^2^ by using the Charpy test procedure. Charpy impact tests were also performed on the samples obtained perpendicular to grains and the results obtained in the current work are shown in Table 14. For the test samples obtained perpendicular to the grains, the maximum impact strength was found to be 0.46 J/cm^2^ and the minimum value was 0.32 J/cm^2^. The average impact strength of Salix Alba Var perpendicular to grains was found out to be 0.38 J/cm^2^.

**Table 13.**
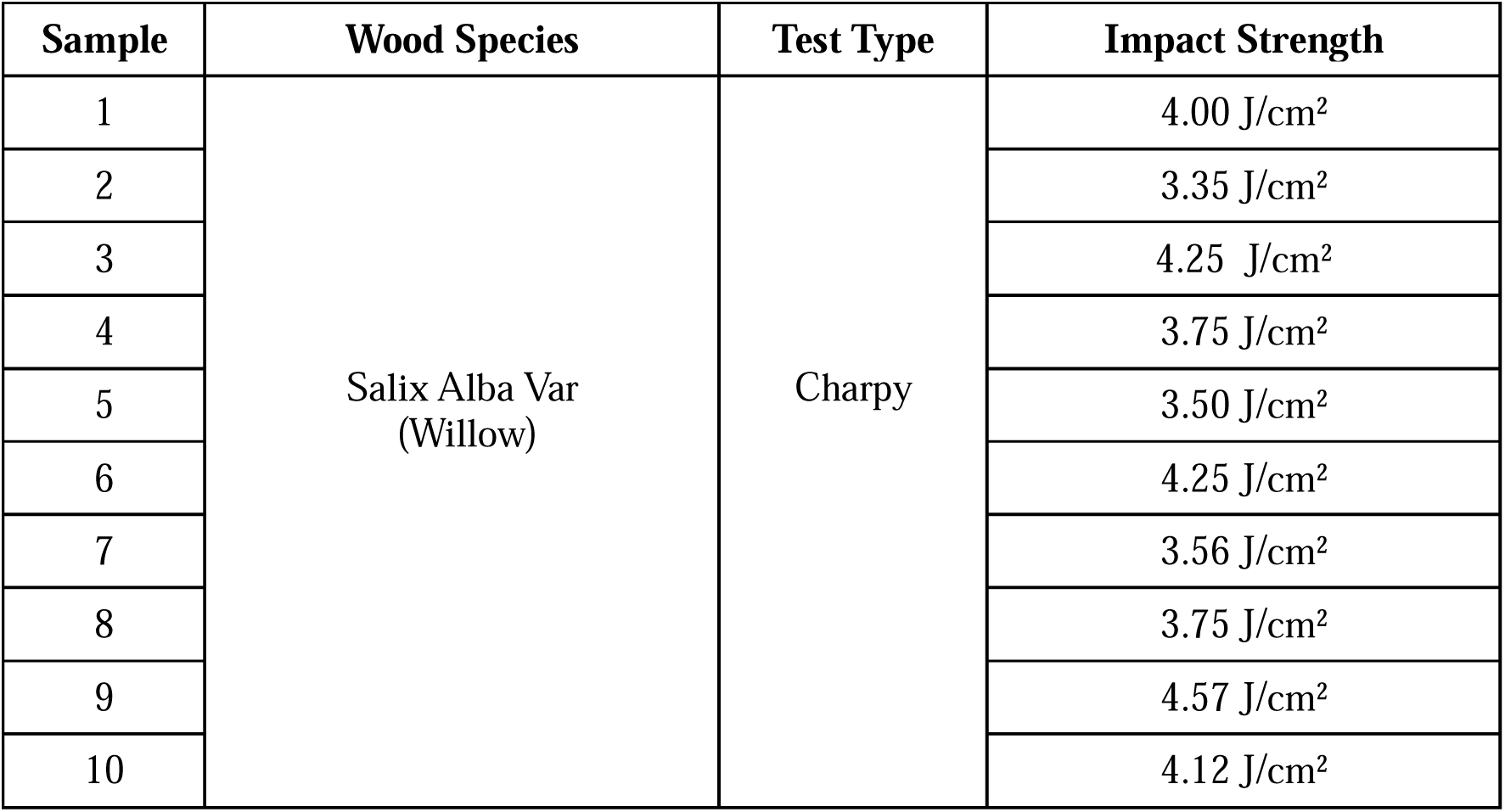
Impact Strength of Salix Alba Var parallel to grains by Charpy test procedure.

**Table 14.** Impact Strength of Salix Alba Var perpendicular to grains by Charpy test procedure.

| Sample | Wood Species | Test Type | Impact Strength |
| --- | --- | --- | --- |
| 1 | Salix Alba Var<br>(Willow) | Charpy | 0.34 J/cm<sup>2</sup> |
| 2 |  |  | 0.33 J/cm<sup>2</sup> |
| 3 |  |  | 0.32 J/cm<sup>2</sup> |
| 4 |  |  | 0.42 J/cm<sup>2</sup> |
| 5 |  |  | 0.36 J/cm<sup>2</sup> |
| 6 |  |  | 0.46 J/cm<sup>2</sup> |
| 7 |  |  | 0.40 J/cm<sup>2</sup> |
| 8 |  |  | 0.34 J/cm<sup>2</sup> |
| 9 |  |  | 0.46 J/cm<sup>2</sup> |
| 10 |  |  | 0.34 J/cm<sup>2</sup> |

### 4.4 Impact Strength of Juglus Riga

Juglus Riga is locally known as “walnut” to the people of Jammu and Kashmir and it is one of the most expensive structural wood available in the region. The testing standards and procedures adopted for determining the impact strength of Juglus Riga are the same as discussed in previous sections. Izod based impact tests were performed on the test specimens obtained from different parts of the northern Himalayan region. The results obtained for the test samples, developed parallel to grains, using Izod impact testing are shown in Table 15. The experimental results show the minimum value of the impact strength as 5.0 J/cm² and the maximum impact strength as 6.0 J/cm². The average impact strength of Juglus Riga in the direction parallel to grains is found to be 5.27 J/cm^2^ by using Izod testing procedure. The impact strength of the test specimens, developed perpendicular to the grains, using Izod impact testing is shown in Table 16. For perpendicular to grain test specimens, the maximum impact strength was found to be 0.61 J/cm² whereas the minimum value was 0.45 J/cm². The average impact strength of Juglus Riga in the direction perpendicular to grains is estimated to be 0.56 J/cm^2^. From the experimental work conducted in the current work, it was observed that Juglus Riga had the highest impact strength as compared to other wood species available in this region.

**Table 15.** Impact strength of Juglus Riga parallel to grains by Izod test procedure.

| Sample | Wood Species | Test Type | Impact Strength |
| --- | --- | --- | --- |
| 1 | Juglus Riga<br>(Walnut) | IZOD | 5.50 J/cm<sup>2</sup> |
| 2 |  |  | 5.00 J/cm<sup>2</sup> |
| 3 |  |  | 6.00 J/cm<sup>2</sup> |
| 4 |  |  | 5.00 J/cm<sup>2</sup> |
| 5 |  |  | 5.18 J/cm<sup>2</sup> |
| 6 |  |  | 5.00 J/cm<sup>2</sup> |
| 7 |  |  | 5.50 J/cm<sup>2</sup> |
| 8 |  |  | 5.00 J/cm<sup>2</sup> |
| 9 |  |  | 5.50 J/cm<sup>2</sup> |
| 10 |  |  | 5.00 J/cm<sup>2</sup> |

**Table 16.** Impact strength of Juglus Riga perpendicular to grains by Izod test procedure.

| Sample | Wood Species | Test Type | Impact Strength |
| --- | --- | --- | --- |
| 1 | Juglus Riga<br>(Walnut) | IZOD | 0.56 J/cm<sup>2</sup> |
| 2 |  |  | 0.60 J/cm<sup>2</sup> |
| 3 |  |  | 0.56 J/cm<sup>2</sup> |
| 4 |  |  | 0.45 J/cm<sup>2</sup> |
| 5 |  |  | 0.55 J/cm<sup>2</sup> |
| 6 |  |  | 0.50 J/cm<sup>2</sup> |
| 7 |  |  | 0.60 J/cm<sup>2</sup> |
| 8 |  |  | 0.61 J/cm<sup>2</sup> |
| 9 |  |  | 0.56 J/cm<sup>2</sup> |
| 10 |  |  | 0.60 J/cm<sup>2</sup> |

Impact strength of Juglus Riga has also been evaluated by using the Charpy impact testing procedure. The impact strength of Juglus Riga parallel to grains by using the Charpy test procedure is given in Table 17. The maximum impact strength parallel to grains was found to be 6.0 J/cm² whereas the minimum impact strength along the same direction was observed as 4.5 J/cm². The average impact strength of Juglus Riga parallel to grains has been evaluated to be 5.23 J/cm² by using the Charpy test procedure. Charpy impact tests were also performed on the samples obtained perpendicular to grains and the results obtained in the current work are shown in Table 18. For the test samples obtained perpendicular to the grains, the maximum impact strength was found to be 0.51 J/cm² and the minimum value was 0.62 J/cm². The average impact strength of Juglus Riga perpendicular to grains was found out to be 0.58 J/cm².

**Table 17.** Impact strength of Juglus Riga parallel to grains by Charpy test procedure.

| Sample | Wood Species | Test Type | Impact Strength |
| --- | --- | --- | --- |
| 1 | Juglus Riga<br>(Walnut) | Charpy | 5.50 J/cm<sup>2</sup> |
| 2 |  |  | 4.50 J/cm<sup>2</sup> |
| 3 |  |  | 6.00 J/cm<sup>2</sup> |
| 4 |  |  | 5.00 J/cm<sup>2</sup> |
| 5 |  |  | 5.18 J/cm<sup>2</sup> |
| 6 |  |  | 4.50 J/cm<sup>2</sup> |
| 7 |  |  | 6.00 J/cm<sup>2</sup> |
| 8 |  |  | 5.50 J/cm<sup>2</sup> |
| 9 |  |  | 5.13 J/cm<sup>2</sup> |
| 10 |  |  | 5.00 J/cm<sup>2</sup> |

**Table 18.** Impact strength of Juglus Riga perpendicular to grains by Izod test procedure.

| Sample | Wood Species | Test Type | Impact Strength |
| --- | --- | --- | --- |
| 1 | Juglus Riga<br>(Walnut) | Charpy | 0.57 J/cm<sup>2</sup> |
| 2 |  |  | 0.61 J/cm<sup>2</sup> |
| 3 |  |  | 0.51 J/cm<sup>2</sup> |
| 4 |  |  | 0.57 J/cm<sup>2</sup> |
| 5 |  |  | 0.55 J/cm<sup>2</sup> |
| 6 |  |  | 0.57 J/cm<sup>2</sup> |
| 7 |  |  | 0.60 J/cm<sup>2</sup> |
| 8 |  |  | 0.61 J/cm<sup>2</sup> |
| 9 |  |  | 0.62 J/cm<sup>2</sup> |
| 10 |  |  | 0.60 J/cm<sup>2</sup> |

### 4.5 Impact Strength of Cedrus Deodara

Cedrus Deodara, locally known as Deodar, is the most widely used structural wood that has found extensive applications in building various engineering structures. The test samples selected for experimentation were defect free and were developed in accordance with the ASTM: D-143 standards. Impact strength of Cedrus Deodara parallel to grains using Izod impact testing procedure is shown in Table 19. The minimum value of the impact strength of this wood species was found to be 2.5 J/cm² whereas the maximum impact strength was 3.5 J/cm². The average impact strength of Cedrus Deodara in the direction parallel to grains is evaluated to be 3.02 J/cm^2^ by using Izod testing procedure. Impact strength has also been evaluated in the direction perpendicular to grains. The impact strength of this wood species perpendicular to the grains using Izod impact testing is shown in Table 20. The maximum impact strength perpendicular to grains was found to be 0.25 J/cm² whereas the minimum value was 0.22 J/cm². The average impact strength of Cedrus Deodara in the direction perpendicular to grains is estimated to be 0.23 J/cm^2^.

**Table 19.** Impact strength of Cedrus Deodara parallel to grains by Izod test procedure.

| Sample | Wood Species | Test Type | Impact Strength |
| --- | --- | --- | --- |
| 1 |  |  | 2.50 J/cm<sup>2</sup> |
| 2 | Cedrus Deodara<br>(Deodar) | IZOD | 3.50 J/cm<sup>2</sup> |
| 3 |  |  | 3.50 J/cm<sup>2</sup> |
| 4 |  |  | 3.75 J/cm<sup>2</sup> |
| 5 |  |  | 2.75 J/cm<sup>2</sup> |
| 6 |  |  | 2.50 J/cm<sup>2</sup> |
| 7 |  |  | 2.75 J/cm<sup>2</sup> |
| 8 |  |  | 3.00 J/cm<sup>2</sup> |
| 9 |  |  | 2.50 J/cm<sup>2</sup> |
| 10 |  |  | 3.50 J/cm<sup>2</sup> |

**Table 20.** Impact strength of Cedrus Deodara perpendicular to grains by Izod test procedure.

| Sample | Wood Species | Test Type | Impact Strength |
| --- | --- | --- | --- |
| 1 | Cedrus Deodara<br>( Deodar) | IZOD | 0.23 J/cm<sup>2</sup> |
| 2 |  |  | 0.22 J/cm<sup>2</sup> |
| 3 |  |  | 0.23 J/cm<sup>2</sup> |
| 4 |  |  | 0.23 J/cm<sup>2</sup> |
| 5 |  |  | 0.25 J/cm<sup>2</sup> |
| 6 |  |  | 0.24 J/cm<sup>2</sup> |
| 7 |  |  | 0.25 J/cm<sup>2</sup> |
| 8 |  |  | 0.23 J/cm<sup>2</sup> |
| 9 |  |  | 0.25 J/cm<sup>2</sup> |
| 10 |  |  | 0.22 J/cm<sup>2</sup> |

In this study, impact strength of Cedrus Deodara has also been evaluated by using the Charpy impact testing procedure. The impact strength of Cedrus Deodara in the direction parallel to grains, by using the Charpy test procedure is given in Table 21. The maximum impact strength parallel to grains was found to be 4 J/cm^2^ whereas the minimum impact strength along the same direction was observed as 2.5 J/cm^2^. The average impact strength of Cedrus Deodara parallel to grains has been evaluated to be 3.22 J/cm^2^ by using the Charpy test procedure. Charpy impact tests were also performed on the samples obtained perpendicular to grains and the results obtained in the current work are shown in Table 22. For the test samples obtained perpendicular to the grains, the maximum impact strength was found to be 0.32 J/cm^2^ and the minimum value was 0.22 J/cm^2^. The average impact strength of Cedrus Deodara perpendicular to grains was found out to be 0.26 J/cm^2^.

**Table 21.** Impact strength of Cedrus Deodara parallel to grains by Charpy test procedure.

| Sample | Wood Species | Test Type | Impact Strength |
| --- | --- | --- | --- |
| 1 |  |  | 3.00 J/cm<sup>2</sup> |
| 2 |  |  | 3.50 J/cm<sup>2</sup> |
| 3 |  |  | 3.50 J/cm<sup>2</sup> |
| 4 | Cedrus Deodara<br>( Deodar) | Charpy | 4.00 J/cm<sup>2</sup> |
| 5 |  |  | 3.00 J/cm<sup>2</sup> |
| 6 |  |  | 2.50 J/cm<sup>2</sup> |
| 7 |  |  | 3.75 J/cm<sup>2</sup> |
| 8 |  |  | 3.00 J/cm<sup>2</sup> |
| 9 |  |  | 2.50 J/cm<sup>2</sup> |
| 10 |  |  | 3.50 J/cm<sup>2</sup> |

**Table 22.** Impact strength of Cedrus Deodara perpendicular to grains by Charpy test procedure.

| Sample | Wood Species | Test Type | Impact Strength |
| --- | --- | --- | --- |
| 1 | Cedrus Deodara<br>( Deodar) | Charpy | 0.23 J/cm<sup>2</sup> |
| 2 |  |  | 0.31 J/cm<sup>2</sup> |
| 3 |  |  | 0.23 J/cm<sup>2</sup> |
| 4 |  |  | 0.31 J/cm<sup>2</sup> |
| 5 |  |  | 0.25 J/cm<sup>2</sup> |
| 6 |  |  | 0.24 J/cm<sup>2</sup> |
| 7 |  |  | 0.25 J/cm<sup>2</sup> |
| 8 |  |  | 0.23 J/cm<sup>2</sup> |
| 9 |  |  | 0.32 J/cm<sup>2</sup> |
| 10 |  |  | 0.22 J/cm<sup>2</sup> |

### 4.6 Impact Strength of Genus Pinus

Genus Pinus is locally known as Pine to the people of Jammu and Kashmir. It has been traditionally used in the making various structures including wooden huts and bridges. The results obtained for the impact strength of Genus Pinus parallel to grains using Izod impact testing procedure are shown in Table 23. The minimum value of the impact strength of this wood species was found to be 2.18 J/cm² whereas the maximum impact strength was 3.5 J/cm². The average impact strength of Genus Pinus in the direction parallel to grains is estimated to be 2.84 J/cm^2^ by using Izod testing procedure. The impact strength of this wood species perpendicular to the grains using the same testing procedure is shown in Table 24. The maximum impact strength perpendicular to grains was found to be 0.28 J/cm² whereas the minimum value was 0.20 J/cm². The average impact strength of Genus Pinus in the direction perpendicular to grains is estimated to be 0.23 J/cm^2^.The impact strength of Genus Pinus in the direction parallel to grains by using the Charpy test procedure is given in Table 25. The maximum impact strength parallel to grains was found to be 3.5 J/cm^2^ whereas the minimum impact strength along the same direction was observed as 2.0 J/cm^2^. The average impact strength of Genus Pinus parallel to grains has been evaluated to be 2.87 J/cm^2^ by using the Charpy test procedure. Charpy impact tests were also performed on the samples obtained perpendicular to grains and the results obtained in the current work are shown in Table 26, where the maximum impact strength was found to be 0.28 J/cm^2^ and the minimum value was 0.15 J/cm^2^. The average impact strength of Genus Pinus perpendicular to grains was found out to be 0.23 J/cm^2^.

**Table 23.** Impact strength of Genus Pinus parallel to grains by Izod test procedure.

| Sample | Wood Species | Test Type | Impact Strength |
| --- | --- | --- | --- |
| 1 | Genus Pinus<br>(Pine) | IZOD | 3.00 J/cm<sup>2</sup> |
| 2 |  |  | 2.50 J/cm<sup>2</sup> |
| 3 |  |  | 2.18 J/cm<sup>2</sup> |
| 4 |  |  | 3.00 J/cm<sup>2</sup> |
| 5 |  |  | 2.75 J/cm<sup>2</sup> |
| 6 |  |  | 2.50 J/cm<sup>2</sup> |
| 7 |  |  | 3.50 J/cm<sup>2</sup> |
| 8 |  |  | 3.00 J/cm<sup>2</sup> |
| 9 |  |  | 2.50 J/cm<sup>2</sup> |
| 10 |  |  | 3.50 J/cm<sup>2</sup> |

**Table 24.** Impact strength of Genus Pinus perpendicular to grains by Izod test procedure.

| Sample | Wood Species | Test Type | Impact Strength |
| --- | --- | --- | --- |
| 1 | Genus Pinus<br>( Pine) | Izod | 0.20 J/cm<sup>2</sup> |
| 2 |  |  | 0.23J/cm<sup>2</sup> |
| 3 |  |  | 0.20 J/cm<sup>2</sup> |
| 4 |  |  | 0.25 J/cm<sup>2</sup> |
| 5 |  |  | 0.28 J/cm<sup>2</sup> |
| 6 |  |  | 0.20 J/cm<sup>2</sup> |
| 7 |  |  | 0.23 J/cm<sup>2</sup> |
| 8 |  |  | 0.20 J/cm<sup>2</sup> |
| 9 |  |  | 0.22 J/cm<sup>2</sup> |
| 10 |  |  | 0.26 J/cm<sup>2</sup> |

**Table 25.** Impact strength of Genus Pinus parallel to grains by Charpy test procedure.

| Sample | Wood Species | Test Type | Impact Strength |
| --- | --- | --- | --- |
| 1 | Genus Pinus<br>( Pine) | Charpy | 2.50 J/cm<sup>2</sup> |
| 2 |  |  | 2.00 J/cm<sup>2</sup> |
| 3 |  |  | 3.00 J/cm<sup>2</sup> |
| 4 |  |  | 3.50 J/cm<sup>2</sup> |
| 5 |  |  | 2.50 J/cm<sup>2</sup> |
| 6 |  |  | 2.50 J/cm<sup>2</sup> |
| 7 |  |  | 3.50 J/cm<sup>2</sup> |
| 8 |  |  | 3.00 J/cm<sup>2</sup> |
| 9 |  |  | 3.50 J/cm<sup>2</sup> |
| 10 |  |  | 2.75 J/cm<sup>2</sup> |

**Table 26.** Impact strength of Genus Pinus perpendicular to grains by Charpy test procedure.

| Sample | Wood Species | Test Type | Impact Strength |
| --- | --- | --- | --- |
| 1 |  |  | 0.15 J/cm<sup>2</sup> |
| 2 |  |  | 0.23 J/cm<sup>2</sup> |
| 3 | Genus Pinus<br>(Pine) | Charpy | 0.28 J/cm<sup>2</sup> |
| 4 |  |  | 0.20 J/cm<sup>2</sup> |
| 5 |  |  | 0.28 J/cm<sup>2</sup> |
| 6 |  |  | 0.19 J/cm<sup>2</sup> |
| 7 |  |  | 0.23 J/cm<sup>2</sup> |
| 8 |  |  | 0.28 J/cm<sup>2</sup> |
| 9 |  |  | 0.20 J/cm<sup>2</sup> |
| 10 |  |  | 0.23 J/cm<sup>2</sup> |

### 4.7 Impact Strength of Abies Pindrow

Abies Pindrow is known as Silver Fir to the local population living in these areas. As discussed earlier, the test samples selected for experimentation were defect free and were developed in accordance with the ASTM: D-143 standards. The results obtained for the impact strength of Abies Pindrow parallel to grains using Izod impact testing procedure are shown in Table 27, where the minimum value of the impact strength can be seen as 1.75 J/cm² whereas the maximum impact strength was 3.0 J/cm². The average impact strength of Abies Pindrow in the direction parallel to grains is estimated to be 2.4 J/cm^2^ by using Izod testing procedure. The impact strength of this wood species perpendicular to the grains using the same testing procedure is shown in Table 28. The maximum impact strength perpendicular to grains was found to be 0.25 J/cm² whereas the minimum value was 0.13 J/cm². The average impact strength of Abies Pindrow in the direction perpendicular to grains is estimated to be 0.19 J/cm^2^.The impact strength of Abies Pindrow in the direction parallel to grains by using the Charpy test procedure is given in Table 29. The maximum impact strength parallel to grains was found to be 3.0 J/cm^2^ whereas the minimum impact strength along the same direction was observed as 1.5 J/cm^2^. The average impact strength of Abies Pindrow parallel to grains has been evaluated to be 2.35 J/cm^2^ by using the Charpy test procedure. Charpy impact tests were also performed on the samples obtained perpendicular to grains and the results obtained in the current work are shown in Table 30, where the maximum impact strength was found to be 0.25 J/cm^2^ and the minimum value was 0.15J/cm^2^. The average impact strength of Abies Pindrow perpendicular to grains was found out to be 0.21 J/cm^2^.

**Table 27.** Impact strength of Abies Pindrow parallel to grains by Izod test procedure.

| Sample | Wood Species | Test Type | Impact Strength |
| --- | --- | --- | --- |
| 1 | Abies Pindrow<br>(Silver Fir) | IZOD | 1.75 J/cm<sup>2</sup> |
| 2 |  |  | 2.50 J/cm<sup>2</sup> |
| 3 |  |  | 1.75 J/cm<sup>2</sup> |
| 4 |  |  | 2.50 J/cm<sup>2</sup> |
| 5 |  |  | 3.00 J/cm<sup>2</sup> |
| 6 |  |  | 2.50 J/cm<sup>2</sup> |
| 7 |  |  | 2.75 J/cm<sup>2</sup> |
| 8 |  |  | 2.00 J/cm<sup>2</sup> |
| 9 |  |  | 2.50 J/cm<sup>2</sup> |
| 10 |  |  | 2.75 J/cm<sup>2</sup> |

**Table 28.** Impact strength of Abies Pindrow perpendicular to grains by Izod test procedure.

| Sample | Wood Species | Test Type | Impact Strength |
| --- | --- | --- | --- |
| 1 | Abies Pindrow<br>(Silver Fir) | IZOD | 0.25 J/cm<sup>2</sup> |
| 2 |  |  | 0.20 J/cm<sup>2</sup> |
| 3 |  |  | 0.18 J/cm<sup>2</sup> |
| 4 |  |  | 0.19 J/cm<sup>2</sup> |
| 5 |  |  | 0.13 J/cm<sup>2</sup> |
| 6 |  |  | 0.25 J/cm<sup>2</sup> |
| 7 |  |  | 0.15 J/cm<sup>2</sup> |
| 8 |  |  | 0.13 J/cm<sup>2</sup> |
| 9 |  |  | 0.15 J/cm<sup>2</sup> |
| 10 |  |  | 0.25 J/cm<sup>2</sup> |

**Table 29.** Impact strength of Abies Pindrow parallel to grains by Charpy test procedure.

| Sample | Wood Species | Test Type | Impact Strength |
| --- | --- | --- | --- |
| 1 | Abies Pindrow<br>(Silver Fir) | Charpy | 2.50 J/cm<sup>2</sup> |
| 2 |  |  | 2.00 J/cm<sup>2</sup> |
| 3 |  |  | 2.28 J/cm<sup>2</sup> |
| 4 |  |  | 2.50 J/cm<sup>2</sup> |
| 5 |  |  | 3.00 J/cm<sup>2</sup> |
| 6 |  |  | 2.50 J/cm<sup>2</sup> |
| 7 |  |  | 1.50 J/cm<sup>2</sup> |
| 8 |  |  | 2.00 J/cm<sup>2</sup> |
| 9 |  |  | 2.50 J/cm<sup>2</sup> |
| 10 |  |  | 2.75 J/cm<sup>2</sup> |

**Table 30.** Impact strength of Abies Pindrow perpendicular to grains by Charpy test procedure.

| Sample | Wood Species | Test Type | Impact Strength |
| --- | --- | --- | --- |
| 1 | Abies Pindrow<br>(Silver Fir) | Charpy | 0.15 J/cm<sup>2</sup> |
| 2 |  |  | 0.23 J/cm<sup>2</sup> |
| 3 |  |  | 0.20 J/cm<sup>2</sup> |
| 4 |  |  | 0.23 J/cm<sup>2</sup> |
| 5 |  |  | 0.20 J/cm<sup>2</sup> |
| 6 |  |  | 0.19 J/cm<sup>2</sup> |
| 7 |  |  | 0.23 J/cm<sup>2</sup> |
| 8 |  |  | 0.25 J/cm<sup>2</sup> |
| 9 |  |  | 0.25 J/cm<sup>2</sup> |
| 10 |  |  | 0.23 J/cm<sup>2</sup> |

### 4.8 Comparative Analysis of All Wood Species

The impact strength of structural wood, obtained from different wood species available in northern Himalayan region of India, has been investigated in the current work. Both Izod and Charpy test procedures were used to evaluate the impact strength of structural wood in both principal directions i.e. parallel and perpendicular to grains. It has been observed that Juglus Riga (Walnut) exhibits maximum impact strength in both principal directions. On the contrary, Abies Pindrow (Silver Fir) showed the minimum impact strength compared to other wood species available in the region. Comparison of the impact strength obtained by the Izod impact testing can be seen in **Figure 14** and **Figure 15**, whereas the results obtained by Charpy test procedures are shown in **Figure 16** and **Figure 17**.

**Figure 14.**
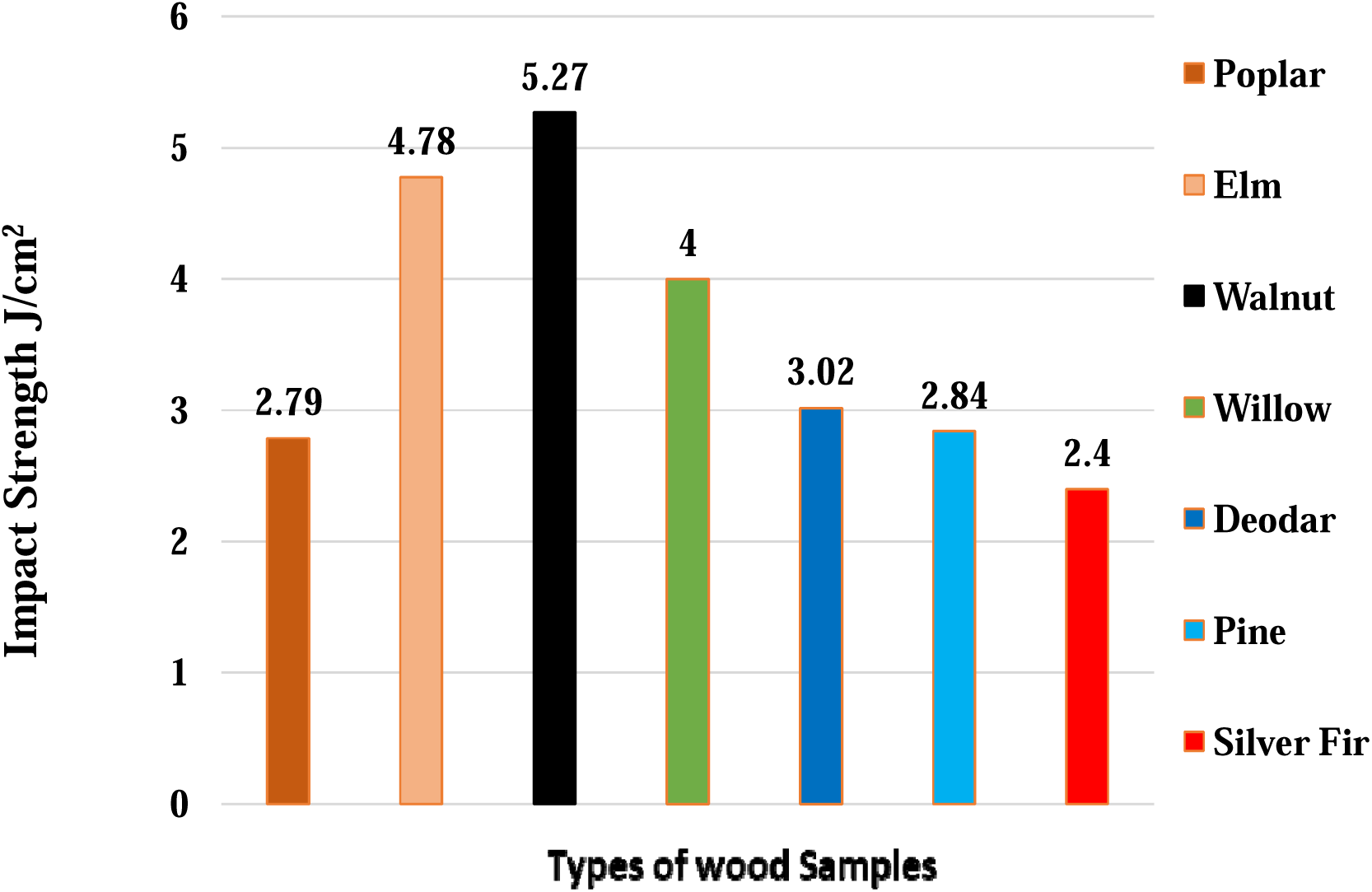
Comparison of average impact strength (parallel to grain) by Izod tests.

**Figure 15.**
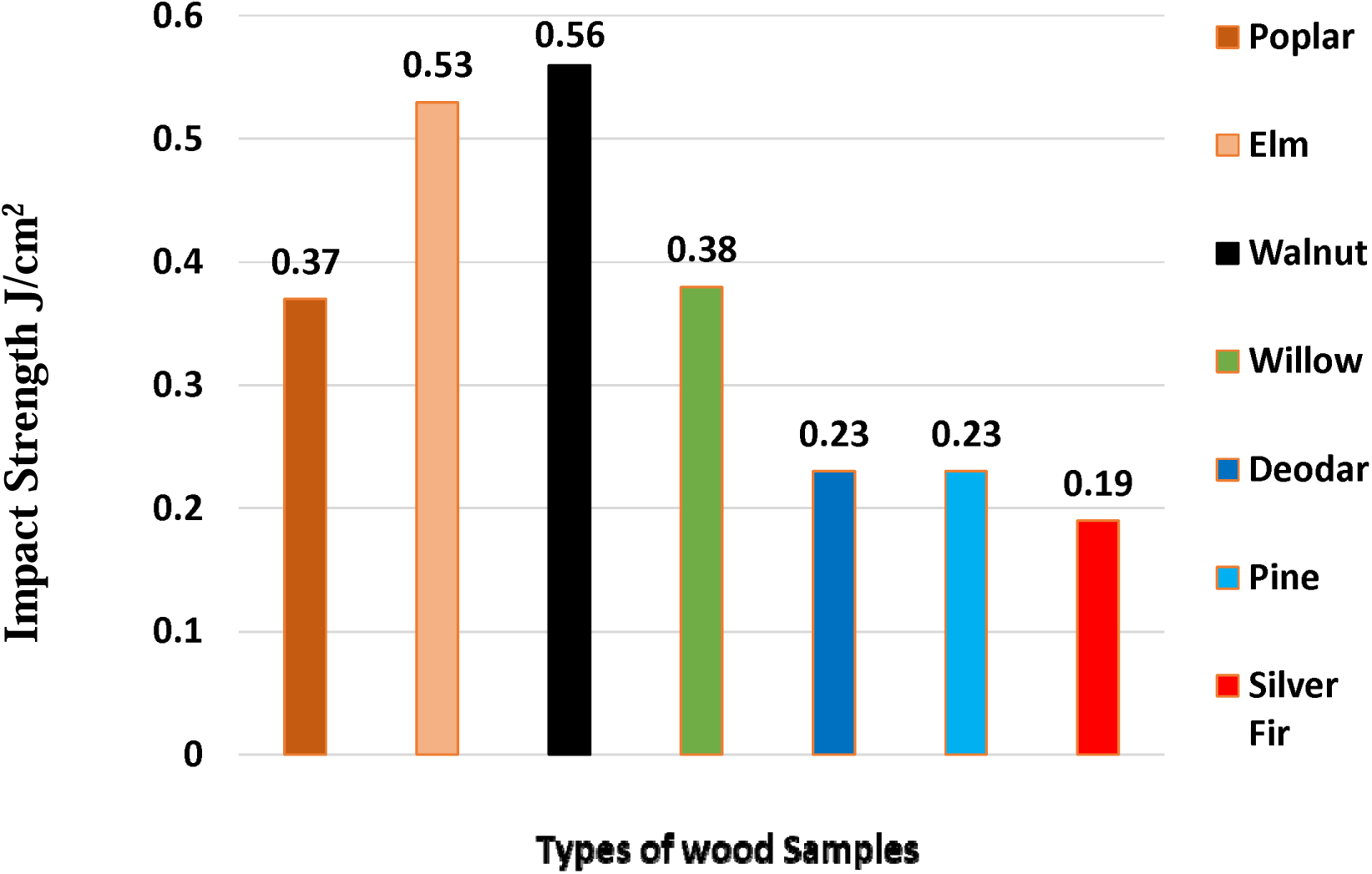
Comparison of average impact strength (perpendicular to grain) by Izod tests

**Figure 16.**
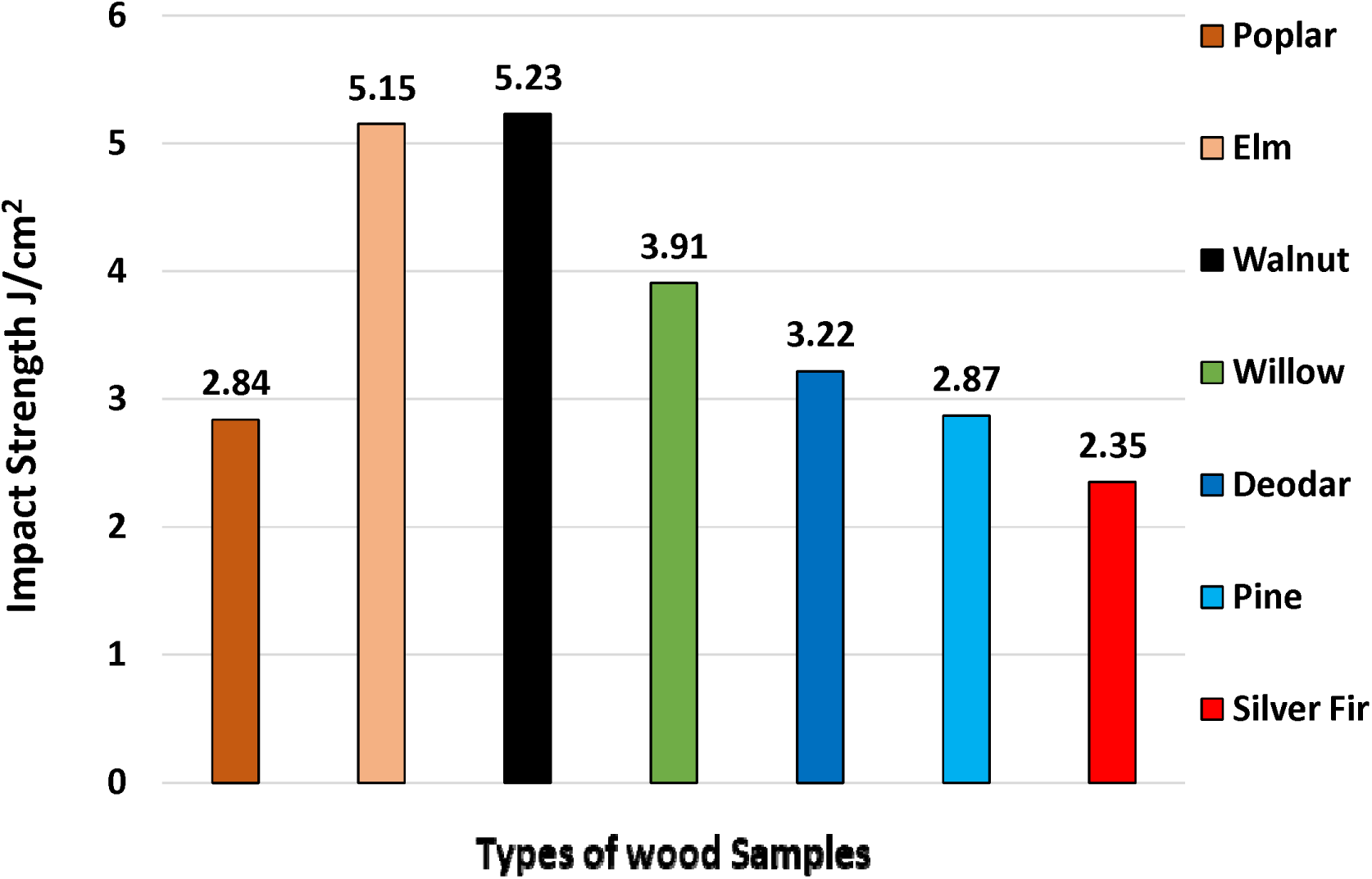
Comparison of average impact strength (parallel to grain) by Charpy tests

**Figure 17.**
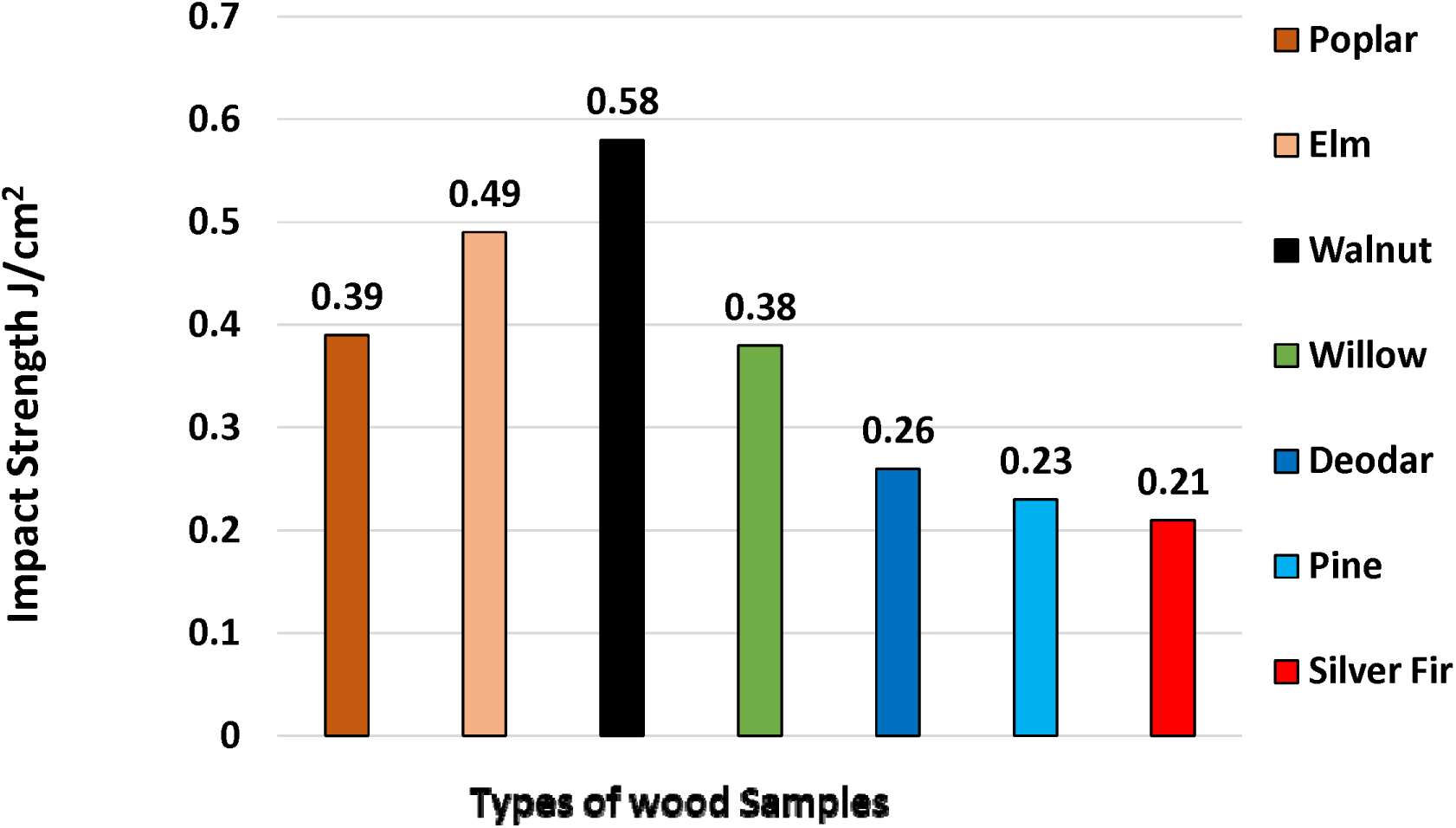
Comparison of average impact strength (perpendicular to grain) by Charpy tests

## Conclusions

The current paper presents the detailed experimental investigations on impact strength of different wood species available in the northern Himalayan region of India. Seven different wood species that have been traditionally used for constructional purposes have been selected for investigation. Defect free test samples were developed in accordance with ASTM: D-143 standards. Wood samples have been seasoned naturally and it has been made sure that the moisture content of the test specimens remains below the fiber saturation point. Both Izod and Charpy test procedures were used to evaluate the impact strength of structural wood in both principal directions i.e. parallel and perpendicular to grains. It has been observed that Juglus Riga (Walnut) exhibits maximum impact strength in both principal directions as compared to other wood species available in the region. On the contrary, Abies Pindrow (Silver Fir) showed the minimum impact strength compared to other wood species available in the region. The experimental data obtained in the current study, regarding the impact strength of structural wood, will play a crucial role in the design of safe and reliable structures in northern Himalayan region of India.

## Notes

### Competing Interest Statement

The authors have declared no competing interest.

